# Neprilysin inhibition reduces microtubule detyrosination in cardiomyocytes through a cGMP-PRKG1-VASH1 axis

**DOI:** 10.64898/2026.03.13.711248

**Authors:** Moritz Meyer-Jens, Chadni Sanyal, Niels Pietsch, Sacnicte Ramirez-Rios, Marisol Herrera-Rivero, Elisabeth Krämer, Ingke Braren, Viacheslav Nikolaev, Maike Frye, Stephanie Könemann, Saskia Schlossarek, Marie-Jo Moutin, Lucie Carrier

**Affiliations:** Institute of Experimental Pharmacology and Toxicology, University Medical Center Hamburg-Eppendorf, Hamburg, Germany; DZHK (German Centre for Cardiovascular Research), partner site North, Hamburg, Germany; University Grenoble Alpes, Inserm, U1216, CNRS, Grenoble Institute of Neurosciences, 38000 Grenoble, France; Department of Cell and Developmental Biology, University of Pennsylvania Perelman School of Medicine, Philadelphia, PA, USA; Department of Genetic Epidemiology, Institute of Human Genetics, University of Münster, Münster, Germany; Institute of Epidemiology and Social Medicine, University of Münster, Münster, Germany; Vector facility, Institute of Experimental Pharmacology and Toxicology, University Medical Center Hamburg-Eppendorf, Hamburg, Germany; Institute of Experimental Cardiovascular Research, University Medical Center Hamburg-Eppendorf, Hamburg, Germany; Institute of Clinical Chemistry and Laboratory Medicine, University Medical Center Hamburg-Eppendorf, Hamburg, Germany; Department of Internal Medicine B, University Medicine Greifswald, Greifswald, Germany; DZHK (German Centre for Cardiovascular Research), partner site North, Greifswald, Germany

## Abstract

Microtubule detyrosination and re-tyrosination on the C-terminus of α-tubulin are mediated by the vasohibin (VASH)-small vasohibin-binding protein (SVBP) complex and tubulin tyrosine ligase (TTL), respectively. Elevated levels of detyrosinated α-tubulin (dTyr-tub) are observed in heart failure, and reducing this modification improves cardiac function, suggesting that clinically used heart failure therapies may modulate microtubule detyrosination. We investigated whether sacubitrilat and valsartan, the active components of the angiotensin receptor–neprilysin inhibitor LCZ696, influence dTyr-tub levels in endothelin-1 (ET1)-induced hypertrophy in human induced pluripotent stem cell-derived cardiomyocytes (hiPSC-CMs). While both sacubitrilat and valsartan prevented hypertrophy, only sacubitrilat prevented ET1-induced dTyr-tub accumulation. RNA sequencing revealed that sacubitrilat normalized several ET1-induced dysregulated pathways. Sacubitrilat slightly increased cyclic guanosine 3’,5’-monophosphate (cGMP) levels and lowered dTyr-tub, whereas inhibition or knockdown of the cGMP-dependent protein kinase 1 (PRKG1) increased dTyr-tub level. Mechanistically, PRKG1 alpha phosphorylated native VASH1. Incubation of microtubules with the VASH1-SVBP complex containing wild-type VASH1 increased detyrosination, while incubation of the complex containing a VASH1 phosphomimic, in which seven C-terminal serine residues were mutated to glutamate (VASH1-7E) did not. Consistently, overexpression of VASH1-7E gave rise to lower dTyr-tub level than overexpression of a non-phosphorylatable form of VASH1 (VASH1-7A) in hiPSC-CMs deficient in VASH1. In conclusion, these findings identify a cGMP–PRKG1–VASH1 signaling axis that reduces microtubule detyrosination in cardiomyocytes. Our work provides mechanistic insight into how neprilysin inhibition may contribute to therapeutic benefit in heart failure.

**One Sentence Summary:** We establish a neprilysin–cGMP–PRKG1–VASH1 signaling axis that reduces microtubule detyrosination in cardiomyocytes.

## INTRODUCTION

Microtubules are part of the cytoskeleton of eukaryotic cells and are composed of α-and β-tubulin subunits that first assemble into heterodimers. Following the exchange of GDP for GTP on β-tubulin, these heterodimers polymerize longitudinally into protofilaments, which then associate laterally to form hollow, cylindrical microtubules. The dynamics and functional properties of microtubules are regulated by various tubulin post-translational modifications (PTMs), including phosphorylation, acetylation, detyrosination, and polyglutamylation (for reviews, see (*1, 2*)). Among these, tubulin detyrosination and re-tyrosination at the C-terminus of α-tubulin were the first PTMs discovered and remain the most extensively studied. Most α-tubulin isoforms are synthesized with a C-terminal tyrosine residue, except for tubulin α8 (TUBA8), which contains a phenylalanine residue and tubulin α4A (TUBA4A), which lacks the C-terminal tyrosine residue. Detyrosination is catalysed by vasohibin 1 (VASH1) and vasohibin 2 (VASH2) in complex with their chaperone, the small vasohibin-binding protein (SVBP; (*3, 4*)) or by the microtubule associated tyrosine carboxypeptidase (MATCAP1; (*5*)). Re-tyrosination is mediated by the tubulin tyrosine ligase (TTL; (*6*)). A key distinction between these enzymes is that TTL acts on free α/β-tubulin dimers, while the VASH-SVBP complex requires polymerised microtubules for binding and subsequent activity. Therefore, modulation of VASH-SVBP binding to microtubules directly affects tubulin detyrosination.

The level of detyrosinated tubulin (dTyr-tub) is consistently elevated across all types of human heart failure (HF), including HF with reduced ejection fraction (HFrEF) or with preserved ejection fraction (HFpEF), dilated cardiomyopathy and hypertrophic cardiomyopathy (HCM; (*7–9*)). Previous studies have shown that reducing dTyr-tub levels either by *VASH1* knockdown or *TTL* overexpression improved contractility in isolated failing human cardiomyocytes (CMs; (*10, 11*)).

Similarly, chronic *TTL* overexpression improved heart function in a mouse model of HCM and normalized cell hypertrophy in HCM human induced pluripotent stem cell (hiPSC)-derived CMs (*12*). Additionally, a small molecule inhibitor of VASH (VASHi) enhanced diastolic function in a rodent model of HFpEF (*13*). The consistent association of elevated dTyr-tub levels in various forms of HF suggests that current state-of-the-art treatments may exert beneficial effects through downstream modulation of microtubule PTMs. Among these, LCZ696 is a dual-acting angiotensin receptor-neprilysin inhibitor, combining the neprilysin inhibitor sacubitril with the angiotensin II receptor blocker valsartan (val), which was approved for the treatment of patients with symptomatic HFrEF (*14*). Previous studies have demonstrated that LCZ696 either prevented or stabilized cardiac dysfunction and reduced fibrosis and hypertrophy in mice subjected to transverse aortic constriction (*15, 16*). Moreover, LCZ696 inhibited angiotensin-II-induced cardiomyocyte hypertrophy in mice (*17*). Our previous study showed that sacubitrilat (sac), the active metabolite of sacubitril, alone or in combination with val prevented the activation of the hypertrophic gene program in a hiPSC-CM model of hypertrophy induced by endothelin-1 (ET1; *16*).

Therefore, the goals of this study were i) to evaluate whether ET1-induced hypertrophy is associated with higher abundance of dTyr-tub and how it is impacted by sac in hiPSC-CMs and ii) to decipher the downstream mechanisms. We show that sac, but not val prevented ET1-induced dTyr-tub in hiPSC-CMs while each compound alone or in combination prevented cellular hypertrophy. We provide evidence that inhibition or knockdown of the cGMP-dependent protein kinase 1 alpha (PRKG1A) reduced dTyr-tub level and that sac moderately increased the cyclic guanosine monophosphate (cGMP) level in hiPSC-CMs. We further show *in vitro* that PRKG1A can phosphorylate native VASH1 and that a phosphomimetic VASH1 (VASH1-7E) in complex with SVBP cannot bind and therefore detyrosinate microtubules. Finally, overexpression of VASH1-7E in VASH1-deficient hiPSC-CMs led to a much lower level of dTyr-tub than overexpression of non-phosphorylatable VASH1 (VASH1-7A). Taken together, this study provides evidence that sacubitrilat reduces microtubule detyrosination in cardiomyocytes through a cGMP-PRKG1-VASH1 axis.

## RESULTS

### Sacubitrilat, but not valsartan, prevents ET1-induced increase in dTyr-tub level in hiPSC-CMs

We used a previously described model of cellular hypertrophy induced by ET1 in hiPSC-CMs (*16, 18*), and evaluated the impact of the individual components of LCZ696, namely sac, val or the combination of both, on cell volume and dTyr-tub level (**Fig. 1A**). ET1 induced a 2.5-fold increase in cell volume of hiPSC-CMs, and all interventions prevented it (**Fig. 1B**). Furthermore, ET1 induced a >1.5-fold increase in dTyr-tub level, which was prevented by the application of sac, but not by val or the combination of both (**Fig. 1C**). These data were supported by the small but significant reduction in dTyr-tub level after an overnight treatment of hiPSC-CMs with sac alone (**Fig. 1D**). These data were further supported by the normalization of the dTyr-tub level after a 6-week treatment with LCZ696 in mice subjected to transverse aortic constriction (*16*) (**Fig. S1A**). Mechanistically, sac inhibits neprilysin, which in turn indirectly increases the abundance of atrial natriuretic peptide (ANP), brain natriuretic peptide (BNP) and C-type natriuretic peptide (CNP) by preventing their degradation. We measured ANP, BNP and CNP concentrations in the medium (**Fig. S1B).** Treatment with sac non-significantly increased the ANP concentration in hiPSC-CM medium from 9 to 11 ng/mL (+23%) but did not impact the level of BNP and CNP (**Fig. S1B**).

**Fig. 1.**
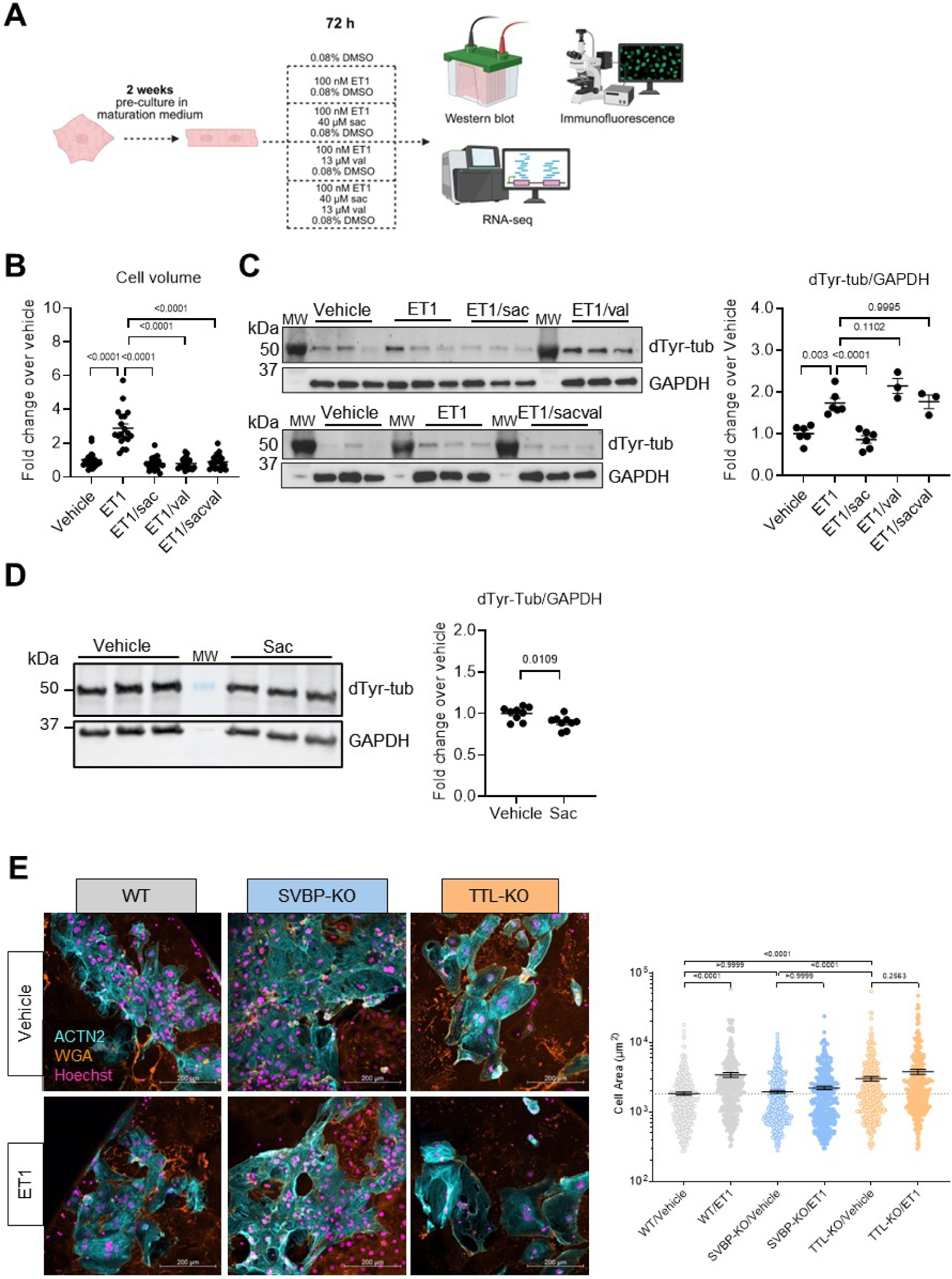
Evaluation of the effect of ET1, sacubitrilat and/or valsartan on cellular hypertrophy and dTyr-tub levels in hiPSC-CMs. (**A**) Protocol: hiPSC-CMs were kept in maturation medium for 2 weeks and then treated for 72 h with vehicle (DMSO, 0.08%), ET1 (100 nM in DMSO), ET1/sac (100 nM/40 µM in DMSO), ET1/val (100 nM/13 µM in DMSO) or ET1/sacval (100 nM/40 µM/13 µM in DMSO); image created with BioRender.com. (**B**) Cell volume measurements of hiPSC-CMs determined by immunofluorescence (N/d= 20/1); 1 outlier (robust regression and outlier removal [ROUT] 1%) was removed from the ET1 group. (**C**) Representative Western Blots of crude hiPSC-CM protein extracts stained for dTyr-tub and GAPDH (loading control) and quantification of dTyr-tub/GAPDH (N/d = 6/2). (**D**) Wild-type hiPSC-CMs were kept in maturation medium for 2 weeks and then treated overnight with vehicle (DMSO, 0.08%) or sac (40 µM in DMSO); representative Western Blot analysis of crude hiPSC-CM protein lysates stained for dTyr-tub and GAPDH (loading control) and quantification of dTyr-tub/GAPDH (N/d=9/2). (**E**) WT, SVBP-KO and TTL-KO hiPSC-CMs were kept for 2 weeks in maturation medium and treated with vehicle (H_2_O) or ET1 (100 nM) for 72 h; representative immunofluorescence images stained for ACTN2 (for cardiomyocytes, blue), WGA (for membrane, orange) and Hoechst (for nuclei, pink) of WT, SVBP-KO and TTL-KO hiPSC-CMs and quantification of cell area (n/N/d=304-382/3/1 each); scale bar, 200 µm. Data are expressed as mean±SEM. Statistical significance was assessed with the unpaired Student’s *t*-test (panels D), with Kruskall-Wallis test and Dunn’s multiple comparisons test (panels B and E) or with one-way ANOVA and Dunnett’s multiple comparisons test vs. ET1 (panel C). Abbreviations: ACTN2, α-actinin 2; dTyr-tub, detyrosinated tubulin; ET1, endothelin-1; hiPSC-CMs, human induced pluripotent stem cell-derived cardiomyocytes; KO, knock-out; MW, molecular weight marker; n/N/d, number of cells/wells/differentiations; sac, sacubitrilat; SVBP, small vasohibin-binding protein; TTL, tubulin tyrosine ligase; val, valsartan; WGA, wheat germ agglutinin; WT, wild-type.

We then tested whether the absence of either VASH-SVBP complex or TTL affects ET1-induced cell hypertrophy. We used previously generated hiPSC lines deficient in either SVBP (SVBP-KO) or TTL (TTL-KO), which developed about 5-fold lower and 19-fold higher dTyr-tub level than wild-type (WT), respectively (*12*). In basal conditions (vehicle), cell area did not differ between SVBP-KO and WT hiPSC-CMs, whereas it was higher in TTL-KO hiPSC-CMs (**Fig. 1E**). ET1 markedly induced hypertrophy in WT, modestly but non-significantly increased cell area in TTL-KO and had no effect in SVBP-KO hiPSC-CMs (**Fig. 1E**). This suggests that the VASH-SVBP detyrosination activity is required for ET1-induced hiPSC-CM hypertrophy.

### Sacubitrilat prevents ET1-induced myofibril assembly and promotes microtubule-based cytoskeleton organization in mitosis

To evaluate molecular changes induced by sac, we performed RNA sequencing (RNA-seq) in WT hiPSC-CMs treated with vehicle, ET1 or ET1/sac. Volcano plots revealed 643 differentially expressed mRNAs (455 higher, 188 lower, log2 ratio >1 or <−1) in ET1- vs. vehicle-treated samples (**Fig. 2A**) and 339 differentially expressed genes (84 higher, 255 lower, log2 ratio >1 or <−1 in ET1/sac- vs. ET1-treated samples (**Fig. 2B**). ET1 induced an increase in expression of genes involved in maturation and hypertrophy, such as *CSRP3* (cysteine and glycine rich protein 3), *DES* (desmin), *MYH7* (myosin heavy chain 7), *NPPB*, *NRAP* (nebulin related anchoring protein) and *XIRP2* (xin actin binding repeat containing 2). All were prevented by the application of sac.

**Fig. 2.**
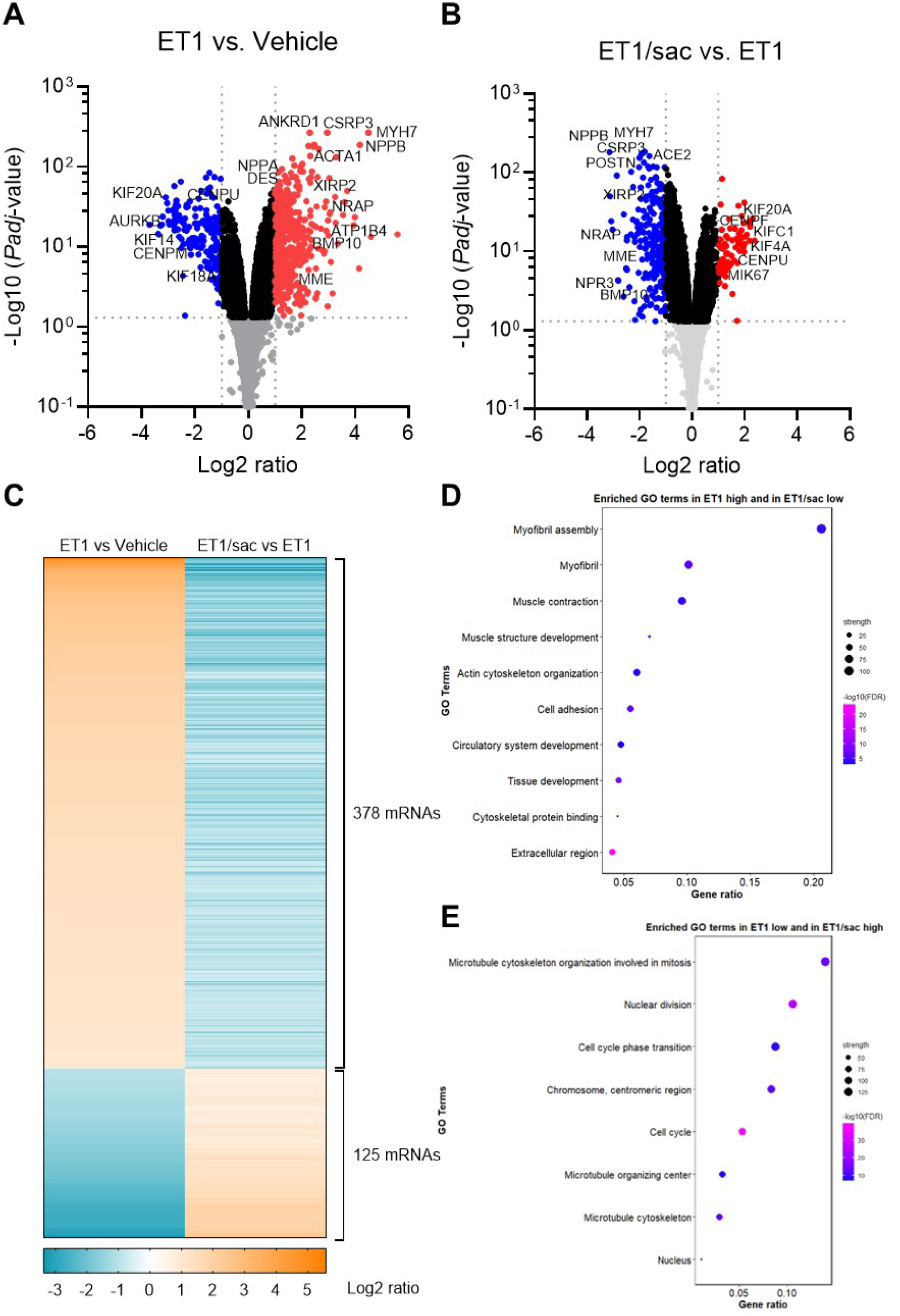
RNA-sequencing analysis in hiPSC-CMs treated with ET1 vs. vehicle or with ET1/sac vs. ET1. RNA-sequencing was performed in wild-type hiPSC-CMs treated with vehicle (DMSO; 0.08%, N=2), ET1 (100 nM in 0.08% DMSO, N=3) or ET1/sac (100 nM/40 µM in 0.08% DMSO, N=3) for 72 h. Volcano plot shows the – log10(P_adj_ value) vs. the magnitude of change (Log2 ratio=log2[case]-log2[reference]) of mRNA levels in ET1-treated vs vehicle-treated hiPSC-CMs **(A)** and in ET1/sac-treated vs. ET1-treated hiPSC-CMs **(B)**. Light gray dots indicate *P*_adj_>0.05, black dots indicate log2 ratio between −1 to +1, red dots indicate the significantly up-regulated genes (Log2 ratio > 1; *P*_Adj_ < 0.05), and blue dots the significantly down-regulated genes (Log2 ratio < 1; *P*_Adj_ < 0.05). **(C)** Heat map of mRNAs which were significantly dysregulated (*P*_adj_<0.05) in both comparisons (ET1 vs. vehicle and ET1/sac vs. ET1); 378 mRNAs exhibited higher level (Log2 ratio >1) in ET1 vs. vehicle and lower level (Log2 ratio < −1) in ET1/sac vs ET1; conversely, 125 mRNAs showed lower level (Log2 ratio < −1) in ET1 vs. vehicle and higher level (Log2 ratio > 1) in ET1/sac vs ET1. **(D)** Top 10 enriched GO terms with a Log2 ratio > 1 in ET1 vs. DMSO and a Log2 ratio < 1 in ET1/sac vs. ET1. **(E)** Top 10 enriched GO terms with a Log2 ratio < 1 in ET1 vs. DMSO and Log2 ratio > 1 in ET1/sac vs. ET1. In panels D,E, the *x* axis shows the gene ratio, the dot size is proportional to the number of gene counts (strength), and the heatmap color shows the extent of *P*_adj_ values (FDR) from the lowest (purple) to the highest (blue). Abbreviations: N, number of samples.

Gene ontology (GO) pathway analysis comparing ET1- vs. vehicle-treated samples revealed an enrichment in components of muscle contraction, extracellular matrix and actin cytoskeleton organization for log2 ratio >1 (**Fig. S2A**) and an enrichment of components of kinetochore organisation, cell division and microtubule cytoskeleton for log2 ratio <−1 (**Fig. S2B**). The opposite pattern was found when we compared ET1/sac- vs. ET1-treated samples with a GO term enrichment in components of cell division and microtubule cytoskeleton for log2 ratio >1 (**Fig. S2C**) and an enrichment in components of heart muscle development and extracellular matrix for log2 ratio <−1 (**Fig. S2D**).

We then extracted only the 503 differentially expressed genes in ET1- vs. vehicle-treated samples that were normalized with sac (378 higher, 125 lower, log2 ratio >1 or <−1; **Fig. 2C; Table S1**). GO term pathway analysis revealed normalization by sac of several components, which were higher expressed in ET1 including myofibril assembly, muscle contraction and actin cytoskeleton organisation (**Fig. 2D**) and lower expressed in ET1 such as microtubule cytoskeleton organisation in mitosis, nuclear division, cell cycle and microtubule cytoskeleton (**Fig. 2E**). By counteracting ET1 effect, sac prevents myofibril assembly and favours microtubule cytoskeleton organization. This suggests a role for downstream pathways of sac in maintaining proliferative capacity during mitosis and limiting hypertrophic differentiation.

### Sacubitrilat activates the cGMP-PRKG1 pathway in hiPSC-CMs

While CNP binds to the natriuretic peptide receptor B (NPRB), both ANP and BNP bind to NPRA. Both receptors contain an intracellular guanylate cyclase domain, which converts GTP to cGMP; cGMP in turn activates PRKG1, which then phosphorylates downstream targets.

We evaluated whether modulation of PRKG1 activity can directly impact the dTyr-tub protein level in hiPSC-CMs. We used RP-8-Br-PET-cGMPS (RP8), which is a cGMP analog acting as a competitive, reversible inhibitor of PRKG1 in hiPSC-CMs (**Fig. 3A**; (*19*)). The dTyr-tub level was 1.4-fold higher in RP8-treated than vehicle-treated cells (**Fig. 3B**). To further investigate if PRKG1 can modulate dTyr-tub level, we performed knockdown experiments with two different commercially available siRNAs directed against *PRKG1* (*siPRKG1*; **Fig. 3A**). The combination of both *siPRKG1* induced a 75% reduction in *PRKG1A* mRNA level (**Fig. 3C**) and a 40% reduction in PRKG1 protein level (**Fig. 3D**) when compared to scrambled (*scr*) siRNA. Simultaneously, *PRKG1* knockdown induced a 25% increase in dTyr-tub level **(Fig. 3E**), supporting the role of PRKG1 in the reduction of microtubule detyrosination.

**Fig. 3.**
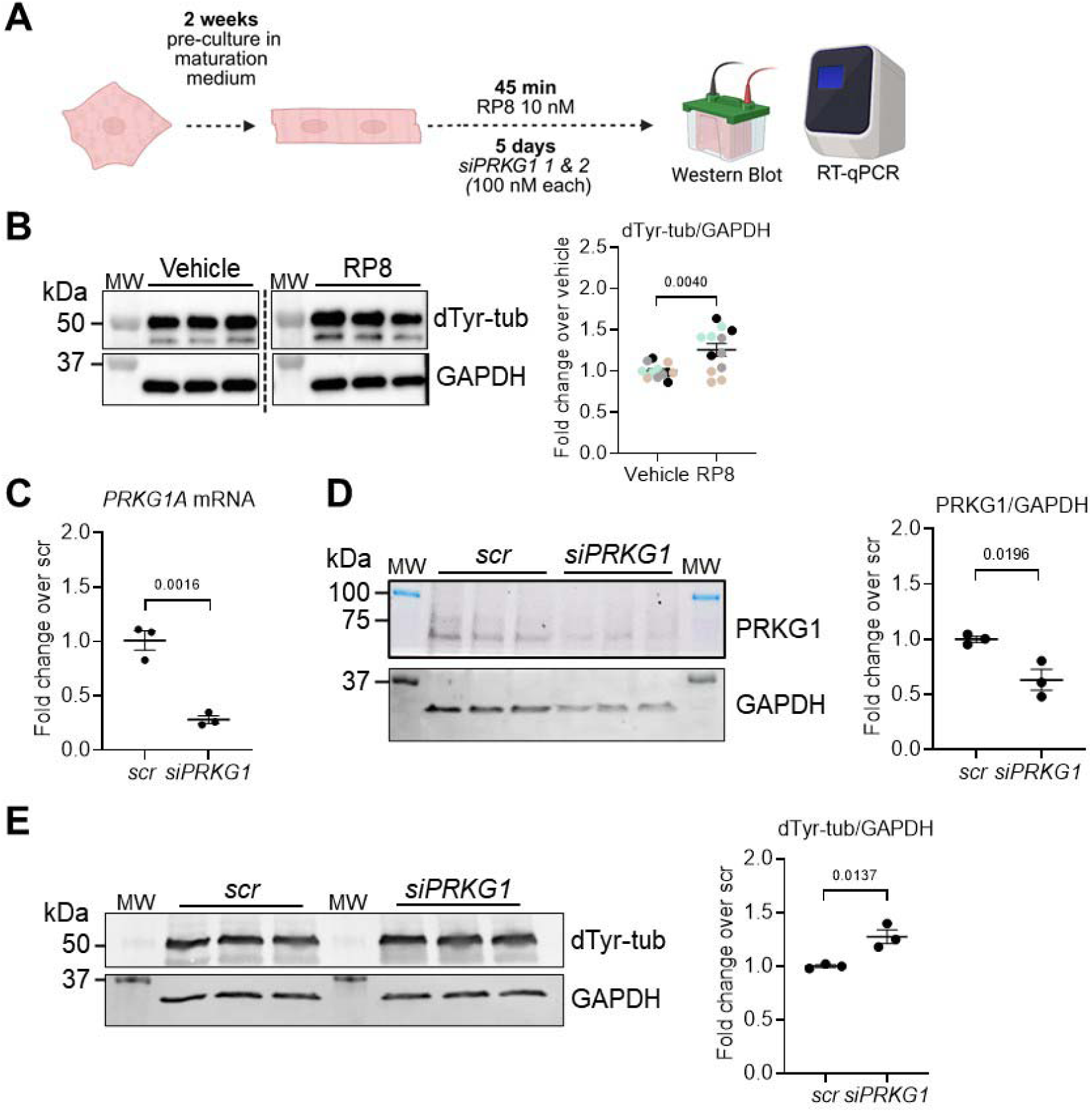
Evaluation of inhibition or knockdown of PRKG1 on dTyr-tub levels in hiPSC-CMs. (**A**) Protocol: hiPSC-CMs were maintained for 2 weeks in maturation medium and subsequently incubated with a PRKG inhibitor (RP8; 10 nM) for 45 min or a combination of 2 *siPRKG1* (100 nM each) for 5 days; image created with BioRender.com. (**B**) Representative Western Blot of crude hiPSC-CM protein lysates stained for dTyr-tub and GAPDH (loading control) and quantification of dTyr-tub/GAPDH after vehicle (H_2_O) or RP8 (10 nM) treatment (N/d=12/4). Colors reflect different batches. (**C**) RT-qPCR for *PRKG1A* to assess the knockdown efficiency after application of 200 nM of *scr* siRNA or 100 nM of 2 different *siPRKG1* (N/d=3/1). (**D**) Western Blot of crude hiPSC-CM protein lysates stained for PRKG1 and GAPDH (loading control) and quantification of PRKG1/GAPDH after application of *scr* siRNA or *siPRKG1* (N/d=3/1). (**E**) Western blot of crude hiPSC-CM protein lysates stained for dTyr-tub and GAPDH (loading control) and quantification of dTyr-tub/GAPDH after application of *scr* siRNA or *siPRKG1* (N/d=3/1). Data are expressed as mean±SEM. Statistical significance was assessed with the unpaired Student’s *t*-test. Abbreviations: dTyr-tub, detyrosinated tubulin; GAPDH, glyceraldehyde-3-phosphate dehydrogenase; hiPSC-CMs, human induced pluripotent stem cell-derived cardiomyocytes; MW, molecular weight marker; N/d, number of wells/differentiations; PRKG1, cGMP-dependent protein kinase 1; RP8, RP-8-Br-PET-cGMPs; scr, scramble; siPRKG1, siRNA directed against PRKG1.

As a positive control for PRKG1 activation, we tested an acute treatment with CNP on a known target of PRKG1, which is the vasodilator-stimulated phosphoprotein (VASP) that is specifically phosphorylated by PRKG1 at serine 239 (pVASP; (*20*)). Treatment with CNP or sac increased the level of pVASP in hiPSC-CMs (**Fig. S3A,B**). We then assessed the intracellular cGMP levels with an enzyme-linked immunosorbent assay (ELISA) following the application of sac or CNP. Sac and CNP increased the cGMP level by about 2- and 8-fold, respectively (**Fig. S3C**). Then, to directly monitor the effect of sac on intracellular cGMP levels, hiPSC-CMs were transduced for 2 days with an adenovirus encoding the cGMP-specific FRET sensor cGi500 (**Fig. S3D**)(*21*). After transduction, the cells were placed in a microscopy chamber suitable for the use with the fluorescence live cell microscope and washed with FRET buffer which was also used to carry out the measurements and dilutions of applied compounds. The application of sac induced about 60% increase in the CFP/YFP ratio (**Fig. S3E**). When the signal started to plateau, a combination of CNP and IBMX (acting as an inhibitor of cAMP and cGMP phosphodiesterases) was added to reach a maximal response (**Fig. S3E**). These results suggest that sac, at least in part, stimulates the intracellular cGMP production downstream of the NPRA and NPRB receptors. Taken together, our data support the implication of the sac-cGMP-PRKG1 pathway in reducing dTyr-tub level in hiPSC-CMs.

### PRKG1A phosphorylates VASH1

To evaluate further down the sac-cGMP-PRKG1 pathway in reducing dTyr-tub in hiPSC-CMs, we investigated whether PRKG1 can phosphorylate VASH1, affect VASH1-SVBP binding on microtubules and therefore reduce microtubule detyrosination *in vitro*. It was previously shown that VASH1-SVBP complex binding to microtubules involves the C-terminal (Ct) domain of VASH1 to promote detyrosination of microtubules (*22*). KinasePhos 2.0 revealed 7 serine residues in this domain (S313, S323, S324, S330, S331, S337, and S344 in human VASH1), which could be phosphorylated by PRKG1 (**Fig. 4A**). We used the previously generated constructs for producing VASH1-SVBP complex containing VASH1-WT or the truncated version of VASH1 containing the core domain (CD) and the Ct domain, consisting of residues 57-365 (VASH1_CD+Ct) (*22*). We also constructed a mutant VASH1 where all 7 serine residues in the Ct domain were mutated to glutamate residues (VASH1-7E; **Fig. 4A**), preventing potential phosphorylation by the kinase.

**Fig. 4.**
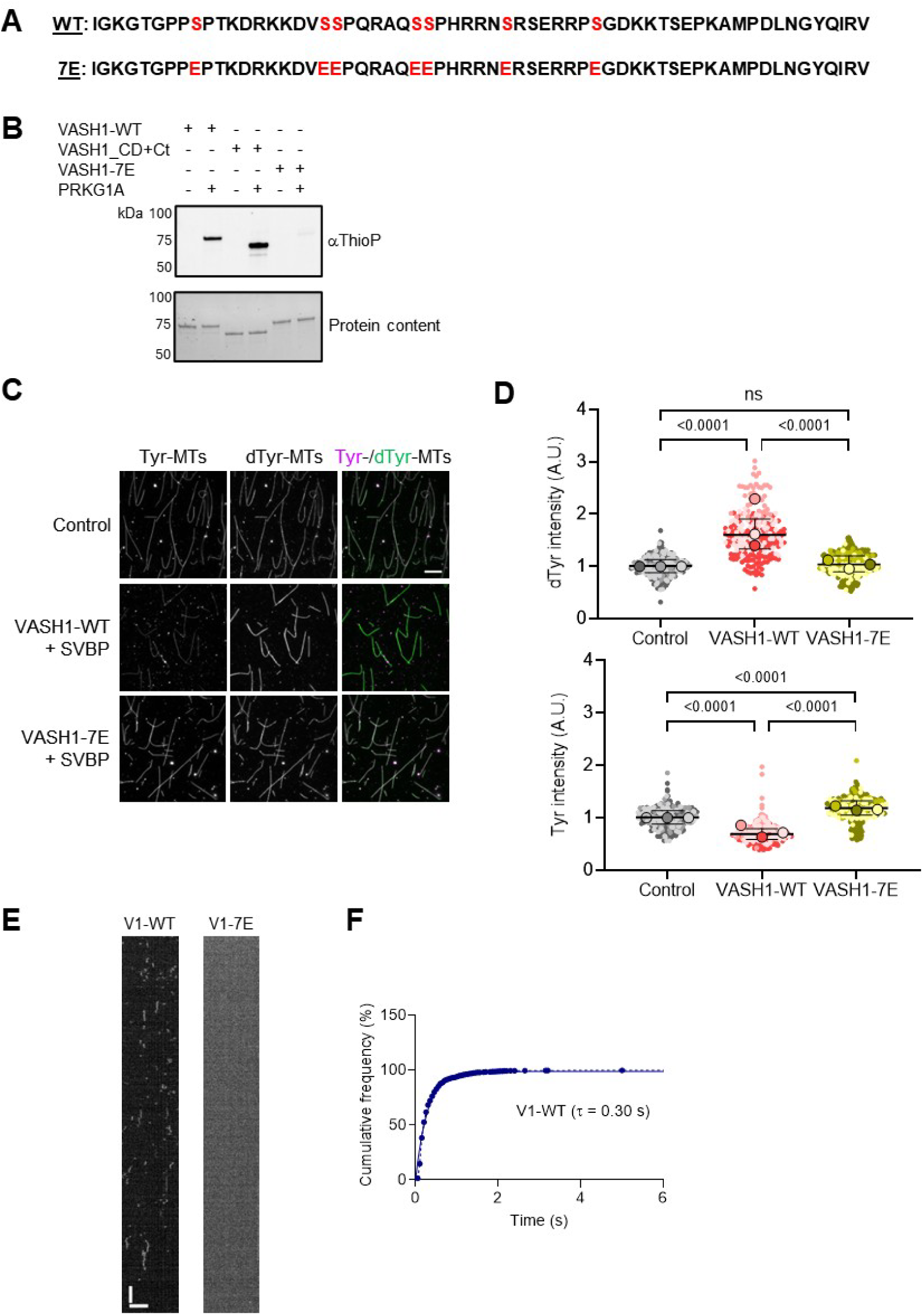
Evaluation of VASH1 as a phosphorylation target of PRKG1A. (**A**) Sequence of the C-terminus of human VASH1-WT and VASH1-7E proteins. The 7 putative serine residues of VASH1-WT which could be phosphorylated by PRKG1A and are mutated to glutamate in VASH1-7E are indicated in red. (**B**) *In vitro* kinase assay of VASH1-SVBP complexes containing VASH1-WT, VASH1_CD+Ct or VASH1-7E proteins in the presence (+) or absence (–) of PRKG1A. Thiophosphorylated proteins were revealed by immunoblots (top) and protein loading by SDS-PAGE (bottom). (**C**) Enzymatic activity of VASH1-WT and VASH1-7E complexes (50 pM) on taxol-stabilized Tyr-MTs. Activity was measured by immunofluorescence in BRB40 supplemented with 50 mM KCl. Representative images of Tyr-MTs (magenta) and dTyr-MTs (green) after 30 min incubation in the absence (control) or presence of the indicated enzymes. Scale bar, 10 µm. (**D**) Quantification of signal intensity of dTyr-MTs (upper panel) and Tyr-MTs (lower panel) in the absence (control) or presence of the indicated enzymes. Each point represents a microtubule (at least 120 microtubules were analyzed per experiment). Data are expressed as median with interquartile range, and as fold change over the control. Colors reflect 3 independent experiments. Statistical significance was assessed with the Kruskal-Wallis test and Dunn’s multiple comparisons test. (**E, F**) Single molecule TIRF microscopy study. (**E**) Representative kymographs of the interaction of single molecules of the sfGFP tagged VASH1 proteins with taxol-stabilized Tyr-MTs in the same experimental conditions as in C-D. Scale bars: horizontal, 5 µm; vertical, 2 min. (**F**) Cumulative frequency of the residence time measured from kymographs. The mean residence time (τ) is obtained by fitting the curve with a mono-exponential function. Abbreviations: A.U., arbitrary unit; CD, core domain; Ct, C-terminus; dTyr-MTs, detyrosinated microtubules; ns, *P* > 0.05; PRKG1A, cGMP-dependent protein kinase 1α; Tyr-MTs, tyrosinated microtubules; VASH1, vasohibin 1; VASH1-7E, VASH1 phosphomimic with 7 glutamate residues; WT, wild-type.

We then performed an *in vitro* non-radioactive kinase reaction in which adenosine 5′-O-(3-thio)triphosphate (ATPγS) served as a phosphate donor. The incorporated ATPγS was alkylated to form thiophosphate ester (αThioP), which was then detected with a specific antibody (*23*). The VASH1-SVBP complex with either VASH1-WT, VASH1_CD+Ct or VASH1-7E was incubated with the vehicle or with PRKG1A and ATPγS. The SDS-PAGE did not show differences in loading in both conditions for the three recombinant proteins, and the subsequent immunoblot stained for αThioP shows a strong signal for VASH1-WT and for VASH1_CD+Ct, whereas only a faint signal was detected for VASH1-7E (**Fig. 4B**). This suggests that PRKG1A phosphorylates at least one of the serine residues present in the Ct domain of VASH1.

We also tested the activity of VASH1-SVBP complexes by immunofluorescence (**Fig. 4C,D**) as described previously (*22*). Taxol-stabilized tyrosinated microtubules were incubated with VASH1-SVBP complex containing either VASH1-WT or VASH1-7E. Incubation with VASH1-WT induced a marked increase in intensity of detyrosination and a marked decrease in intensity of tyrosination of microtubules respectively. In contrast, incubation with the phosphomimetic VASH1-7E did not increase the intensity of microtubule detyrosination and induced a significantly higher intensity of microtubule tyrosination than VASH1-WT and control.

Additionally, we monitored the interaction behaviour of VASH1-WT and VASH1-7E with the microtubules using single molecule Total Internal Reflection Fluorescence (TIRF) microscopy. Representative kymographs show the expected binding pattern of VASH1-WT to the microtubules (short and frequent binding events) but no binding of VASH1-7E (**Fig. 4E**). The average residence time of VASH1-WT over the 30 min of incubation with the tyrosinated microtubules was 0.3 s with a monoexponential distribution (**Fig. 4F**), which is in range with previous findings in similar experimental conditions (*22*). Together, these data provide evidence that adding negatively charged residues at potential phosphorylation sites in the Ct domain of VASH1, i.e. mimicking the phosphorylated state, affects binding of the VASH1-SVBP complex and thereby its detyrosination activity on microtubules.

### VASH1 phosphomimic has a lower detyrosination activity than non-phosphorylatable VASH1 in hiPSC-CMs

We then evaluated whether the VASH1-SVBP complex containing VASH1-7E affects dTyr-tub levels in hiPSC-CMs. We created and characterized a VASH1-KO hiPSC line with CRISPR/Cas9 genetic tools (**Fig. S4A-C**). VASH1-KO hiPSCs were differentiated into CMs, and the level of dTyr-tub normalized to total α-tubulin was 65% lower in VASH1-KO than in WT hiPSC-CMs, respectively (**Fig. S4D-F**). The remaining basal level of dTyr-tub in VASH1-KO hiPSC-CMs could be mediated by other enzymes, such as VASH2-SVBP and MATCAP1, or by TUBA4A, an α-tubulin encoded without the C-terminal tyrosine. We generated lentiviruses encoding YFP-tagged mouse VASH1+SVBP containing either VASH1-WT, the phosphomimetic VASH1-7E or a non-phosphorylatable VASH1, where all 7 serine residues of the Ct domain of VASH1 were mutated to alanine residues (VASH1-7A). We overexpressed the three constructs in hiPSC-CMs deficient in VASH1 and analyzed the levels of VASH1-YFP, dTyr-tub and Tyr-tub (**Fig. 5A**). The level of exogenous VASH1-YFP was >60% lower in hiPSC-CMs transduced with VASH1-7E or VASH1-7A than with VASH1-WT (**Fig. 5B**). The level of dTyr-tub was 22-fold higher and Tyr-tub level was 50% lower in VASH1-WT-expressing cells than in untreated VASH1-KO cells (**Fig. 5C**). Comparing directly the overexpression of VASH1-7E and VASH1-7A, we did not see differences in the exogenous level of YFP-VASH1 between the two groups (**Fig. 5D**), and dTyr-tub level was 2.6-fold lower and Tyr-tub level 11% higher in cells expressing VASH1-7E than in cells expressing VASH1-7A (**Fig. 5D**). Taken together, these cellular data show that mimicking phosphorylation by adding negative charges on VASH1 C-terminal serine residues reduces microtubule detyrosination.

**Fig. 5.**
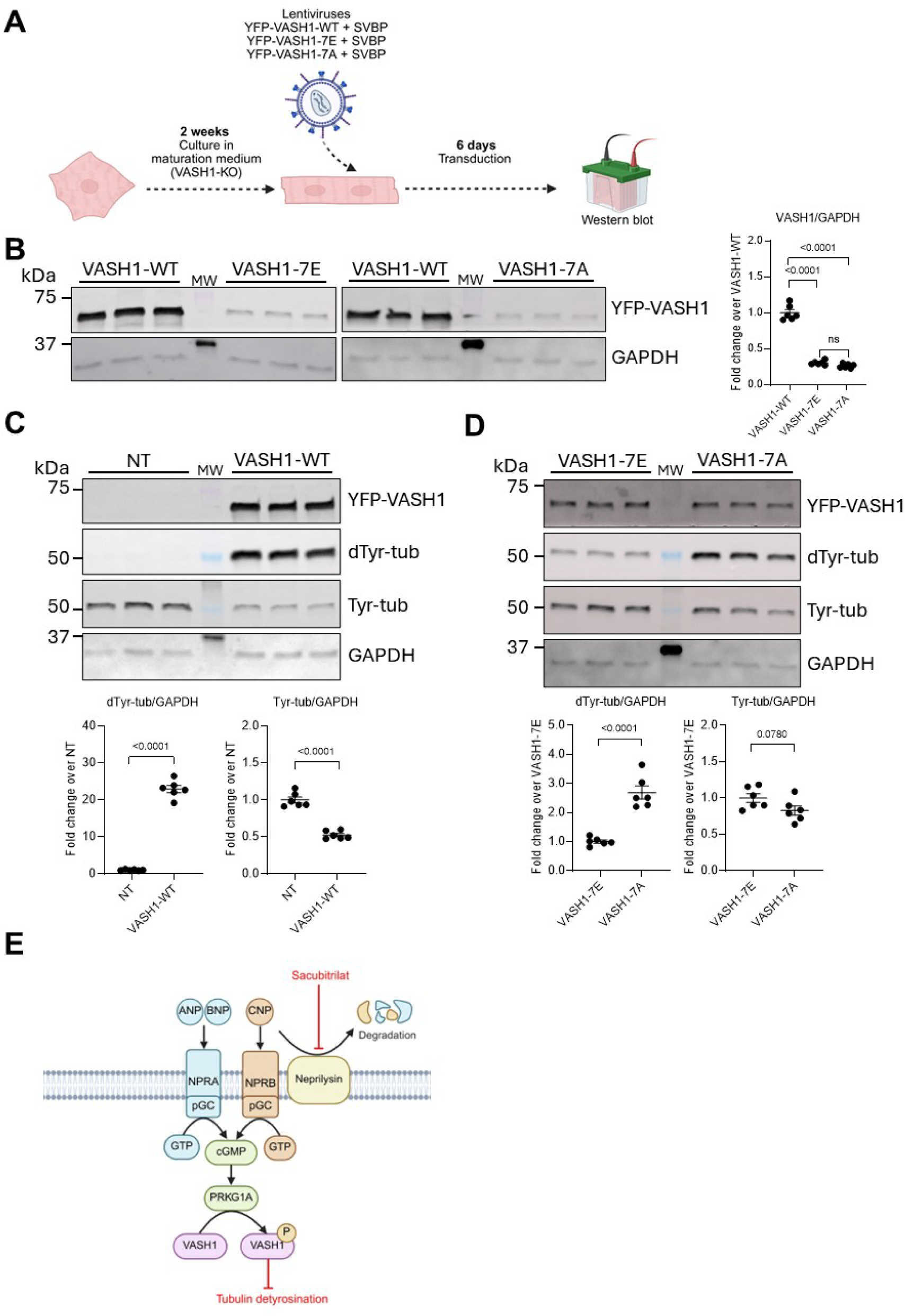
Evaluation of the overexpression of VASH1-WT, VASH1-7E and VASH1-7A on tubulin modifications in VASH1-KO hiPSC-CMs. **(A)** Protocol: VASH1-KO hiPSC-CMs were kept in maturation medium for 2 weeks and then transduced for 6 days with lentivirus expressing YFP-tagged VASH1-WT, VASH1-7E or VASH1-7A each in combination with SVBP. **(B)** Representative Western blot of crude hiPSC-CM protein lysates stained for YFP-VASH1 and GAPDH (loading control) and quantification of VASH1/GAPDH in the 3 conditions (N/d=6/1). **(C)** Representative Western blot of crude hiPSC-CM protein lysates stained for YFP-VASH1, dTyr-tub, Tyr-tub and GAPDH (loading control) and quantification of dTyr-tub/GAPDH and Tyr-tub/GAPDH in non-transduced VASH1-KO hiPSC-CMs or transduced with YFP-VASH1-WT in combination with SVBP (N/d=6/1). **(D)** Representative Western blot of crude hiPSC-CM protein lysates stained for YFP-VASH1, dTyr-tub, Tyr-tub and GAPDH (loading control) and quantification of dTyr-tub/GAPDH and Tyr-tub/GAPDH in VASH1-KO hiPSC-CMs transduced with YFP-VASH1-7E or YFP-VASH1-7A in combination with SVBP (N/d=6/1). **(E)** Proposed model of regulation of VASH1 activity downstream of the action of sac. Sac inhibits neprilysin that would otherwise degrade the natriuretic peptides ANP, BNP and CNP. Through this inhibition, these natriuretic peptides can bind their respective receptors, which leads to the intracellular production of cGMP. Increased amounts of intracellular cGMP would then activate PRKG1A and thereby phosphorylate different targets, including VASH1. Eventually, phosphorylation of VASH1 reduces the binding of VASH1 to microtubules and tubulin detyrosination. Data are expressed as mean±SEM. Statistical significance was assessed with the one-way ANOVA and Tukey’s multiple comparisons tests (panel B) or unpaired Student’s *t*-test (panels C and D). Abbreviations: ANP, atrial natriuretic peptide; BNP, brain natriuretic peptide; cGMP, guanosine 3′,5′-cyclic monophosphate; CNP, C-type natriuretic peptide; dTyr-tub, detyrosinated tubulin; GAPDH, glyceraldehyde-3-phosphate dehydrogenase; GTP, guanosine triphosphate; hiPSC-CMs, human induced pluripotent stem cell-derived cardiomyocytes; KO, knock-out; MW, molecular weight marker; N/d, number of wells/differentiations; NPRA, natriuretic peptide receptor 1; NPRB, natriuretic peptide receptor 2; P, phosphorylated; pGC, particulate guanylate cyclase; PRKG1A, cGMP-dependent protein kinase G1 alpha; Tyr-tub, tyrosinated tubulin; VASH1, vasohibin 1; VASH1-7A, non-phosphorylatable VASH1 with 7 alanine residues; VASH1-7E, VASH1 phosphomimic with 7 glutamate residues; WT, wild-type.

We therefore propose that sacubitrilat reduces tubulin detyrosination in cardiomyocytes through a cGMP-PRKG1-VASH1 axis (**Fig. 5E**).

## DISCUSSION

In this study we evaluated the effect of sacubitrilat and valsartan, the active components of the angiotensin receptor-neprilysin inhibitor LCZ696 on α-tubulin detyrosination in a human model of cardiomyocyte hypertrophy induced by endothelin-1. We provide evidence that the activation of the cGMP/PRKG1 pathway by sacubitrilat promotes phosphorylation of VASH1 and reduces dTyr-tub levels in hiPSC-CMs.

We show that ET1 induced cellular hypertrophy in WT, but not in SVBP-KO, and did not increase further the already higher cell area in TTL-KO hiPSC-CMs. This suggests a central role for the VASH-SVBP complex – and therefore increased microtubule detyrosination in the development of hypertrophy. Furthermore, RNA-seq data analysis of hiPSC-CMs treated with ET1 revealed an enrichment in pathways related to myofibril assembly and organisation of actin cytoskeleton, with specific accumulation of transcripts involved in hypertrophy, such as *CSRP3*, *DES, MYH7*, *NPPA*, *NPPB*, *XIRP2*, and on the contrary, reduction in levels of transcripts related to cell cycle (such as aurora kinase B, centromere protein U, proliferation marker protein Ki-67) and microtubule organisation (such as kinesin family member 14 and 20A). These findings support previous RNA-seq data obtained in hiPSC-CMs under similar conditions (*18*). We show that the addition of sacubitrilat to ET1 prevented all these changes, suggesting not only a broad antihypertrophic effect but also a potential pro-proliferative effect of sacubitrilat. The beneficial effect of sacubitrilat might in part be due to preventing the upregulation of *CSRP3* (*24*). These authors indeed reported that the NO donor S-nitroso-N-acetylpenicillamine (SNAP) significantly suppressed ET1-induced *CSRP3* expression and cardiomyocyte hypertrophy (*24*). Likewise, *XIRP2* is not only implicated in the maturation of cardiomyocytes (*25*) but also plays a role in cardiomyocyte hypertrophy (*26*) and as a gene modifier for HCM (*27*). *XIRP2* transcript level was also elevated in *Mybpc3*-knockout mice, indicating a contribution to the pathophysiological remodelling in these mice (*28*). Another study reported a similar effect of sacubitrilat/valsartan or valsartan alone on the reduction of transcripts levels of *Myh7* and *Acta1* in Langendorff-perfused mouse hearts while also outlining the superior effect of sacubitrilat/valsartan even over an equimolar dose of valsartan alone (*15*). The effect of sacubitrilat/valsartan was also superior to valsartan on cardiac fibrosis, which they attributed to a restoration of normal PRKG1 signaling in cardiac fibroblasts. Other groups have also shown that desmin is required for the stabilization of growing microtubules alongside with dTyr-tub (*29*), making it a likely contributor in driving hiPSC-CM hypertrophy after ET1 exposure. Normalization of *DES* mRNA level by additional sacubitrilat treatment is interesting especially when considering the effects on dTyr-tub we report here.

In our ET1-based hiPSC-CM hypertrophy model, all treatments effectively prevented the increase in cell volume. The effect of valsartan on preventing cellular hypertrophy aligns with its established role as an angiotensin-II receptor antagonist (*30*). Previous work has shown that sacubitrilat and valsartan have differential roles in mouse models of heart failure with preserved ejection fraction with the combinational treatment being superior compared to valsartan alone (*31*). Specifically, this latter study reported a larger decrease in myocyte cross-sectional area and a better improvement of diastolic function with the combination of sacubitrilat and valsartan than with valsartan alone. They attribute these effects to a potential increase in cGMP levels, which serves as an activator for PRKG1.

Our study shows that the angiotensin receptor-neprilysin inhibitor LCZ696 normalized dTyr-tub level in mice subjected to transverse aortic constriction, whereas the combination of sacubitrilat and valsartan did not in hiPSC-CMs treated with ET1. On the contrary, sacubitrilat prevented the ET1-induced increase in dTyr-tub level in hiPSC-CMs. These data suggest that the effect of sacubitrilat is mediated by the inhibition of neprilysin and reduced degradation of natriuretic peptides. By contrast, the lack of effect of the combination treatment on dTyr-tubulin levels remains unclear; one possible explanation is that valsartan masked the effect of sacubitrilat under ET1 conditions. To date, clinical studies assessing the impact of LCZ696 or its individual components on dTyr-tub levels are lacking. However, previous studies have shown that an elevated level of dTyr-tub contributes to impaired cardiac function in several types of heart failure (*10, 11, 32*). We also provide evidence that a treatment with sacubitrilat alone reduced dTyr-tub level, tended to increase ANP release in the medium of hiPSC-CMs and increased the intracellular cGMP concentration, indicating that it can modulate cGMP-signalling in hiPSC-CMs.

Previous studies have shown that overexpression of *TTL*, knockdown of *VASH1* or inhibition of VASH1/2 activity with EpoY derivatives effectively reduces dTyr-tub level and restores contractility in isolated human failing cardiomyocytes and *in vivo* in animal models (*10–13*). These improvements were primarily attributed to decreased tissue or cellular stiffness (*7*). Notably, knockdown of *VASH2* was less efficient than of *VASH1* in reducing dTyr-tub level, and VASH1 is the major cardiac isoform (*10*), suggesting VASH1 as a relevant therapeutic target for heart failure.

Moreover, we show that a reduction of PRKG1 abundance with siRNA or PRKG1 activity with the RP8 small molecule inhibitor (*33*) increased dTyr-tub by 25% and 50%, respectively. PRKG1 inhibitors have often been reported to have suboptimal target specificity (*34*), including RP8 (*35*), which can be seen as a limitation of our study. On the other hand, hiPSC-CMs mainly express *PRGK1A*, suggesting that the applied treatments can only work on PRKG1A in hiPSC-CMs. It would be of interest to test more specific PRKG1 inhibitors to substantiate the generated data, but these are lacking. Taken together, these data indicate that the cGMP-PRKG1 pathway is active in hiPSC-CMs and its stimulation reduces dTyr-tub level.

We provide evidence that PRKG1A phosphorylates VASH1 *in vitro.* Furthermore, we demonstrated that the VASH1-SVBP complex containing unphosphorylated VASH1 binds to microtubules and increases dTyr intensity, whereas the complex containing the phosphomimetic VASH1-7E does not. We also showed that VASH1-7E overexpression resulted in a lower amount of dTyr-tub than the non-phosphorylatable VASH1-7A in VASH1-KO hiPSC-CMs. The phosphorylation sites of PRGK1A on VASH1 were found to be in the C-terminus of the protein. This section of the VASH protein has been shown to be important for the binding of both VASH1 and VASH2 to microtubules and is rich in basic amino acids while the N-termini differ and are important for differentiating their function and activity (*22*). Previous research has elucidated that phosphorylation can disrupt enzyme-target binding by several mechanisms such as steric hinderance, electrostatic repulsion, motif disruption and conformational changes (*36–38*). It is likely that changed electrostatic properties of the otherwise basic VASH1 C-terminus by phosphorylation impedes binding to the acidic surface of microtubules (*39*). As VASH1 can only perform its carboxypeptidase activity when bound to microtubules (*3, 40*), phosphorylation by PRKG1A likely represents a negative regulatory mechanism for the enzyme.

In conclusion, this study shows that sacubitrilat reduces microtubule detyrosination in cardiomyocytes through a cGMP-PRKG1-VASH1 axis. This may contribute to the therapeutic benefit of LCZ696 in patients with heart failure.

## MATERIAL AND METHODS

### 2D culture of hiPSC-CMs

The WT hiPSC line mTagRFPT-TUBA1B (AICS-0031-035, Coriell Institute, Camdem, NJ, US) was utilized as a control line for this study. Additionally, we used SVBP-KO and TTL-KO hiPSC lines, that were generated and characterized in our lab from the above-mentioned isogenic control line (*12*). The creation and characterization of VASH1-KO hiPSC line is given in **Fig. S4**. In brief, two gRNAs were utilized to excise a 16,144 bp fragment from the VASH1 locus including almost the entire coding sequence of VASH1 (**Fig. S4A**). HiPSCs were analyzed by PCR for successful editing (**Fig. S4B**) and quality controlled by NanoString™ karyotype profiling (**Fig. S4C**), off-target analysis of the ten most likely off-target loci per gRNA (not shown) and tested for the absence of mycoplasma (not shown). VASH1-KO could be differentiated into hiPSC-cardiomyocytes with high purity (**Fig. S4D**) and showed strongly reduced levels of dTyr-tub (**Fig. S4F**). HiPSC were maintained in in-house medium (DMEM F-12, L-glutamine 2 mM, transferrin 5 µg/mL, selenium 5 ng/mL, human serum albumin 0.1%, lipid mixture 1X, insulin 5 µg/mL, dorsomorphin 50 nM, activin A 2.5 ng/mL, TGFβ1 0.5 ng/mL, bFGF 30 ng/mL) while being seeded at an initial density of 6.5 × 10^4^ hiPSCs/cm^2^. Cells were fed daily, kept under hypoxia and monitored microscopically for culture density and morphology. Passaging was performed at 90% confluency with the medium being supplemented with 10 µM Y-27632. The differentiation of hiPSCs to CMs was performed by adapting a monolayer protocol from (*41*). The day before starting the differentiation protocol, cells were passaged onto 6-well plates coated with Corning Matrigel (354234, high growth factor) at 5.2 × 10^4^ hiPSC/cm^2^. The following day, cells were fed in the morning, and the regular culture medium was replaced in the evening with stage 0 medium (StemPro-34 SFM, StemPro Supplement 2.6%, BMP4 1 ng/mL, L-glutamine 2 mM, matrigel high growth factor 1%). Twelve to 16 h later, stage 0 was removed and stage 1 (StemPro-34 SFM, StemPro supplement 2.6%, BMP4 10 ng/mL, L-glutamine 2 mM, activin A 8 ng/mL) was added for the specification of cardiac progenitor cells. In subsequent 48 h steps, stage 2.1 (RPMI1640, B27 2%, KYO2111 10 µM, XAV939 10 µM) and 2.2 (stage 2.1 with insulin 3.21 µg/mL) were used to inhibit Wnt signaling. Two days after stage 2.2 cells were switched to feeding medium (RPMI1640, B27 5%, insulin 3.21 µg/mL). This feeding schedule was continued until the cells started beating homogenously. Cells were dissociated as described before (*16*) and plated at the indicated densities onto Geltrex (Gibco) coated 96-/12-/6-well plates. Three days after plating the complete medium (DMEM, Sigma, D5546; 10% FCS, Invitrogen, 26140079; 1% Pen-Strep, Gibco, 15140122; 10 µg/mL Insulin, Sigma, I9278) was exchanged for maturation medium (*42*); DMEM w/o glucose, glucose 3 mM, L-lactate 10 mM, vitamin B12 5 µg/mL, biotin 0.82 µM, creatine monohydrate 5 mM, taurine 2 mM, L-carnitine 2 mM, ascorbic acid 0.5 mM, NEAA 1X, albumax I 0.5 %, B27 1X, knockout serum replacement 1X, penicillin-streptomycin 1%, insulin 10 µg/mL). Cells were kept in this medium for 2 weeks with medium being changed every five days. Cells were treated with 100 nM ET1 (Sigma Aldrich, 05-23-3800) alone or in combination with 40 µM sac (Sigma Aldrich, SML2064), 100 nM CNP (Tocris, 3520), or varying concentrations of ANP (Tocris, 1906), or RP8 (Tocris, 3028) at indicated concentrations. Sac, ET1/sac, ET1/val, ET1/sacval, were dissolved in DMSO, while ANP, CNP, ET1 and RP8 were dissolved in sterile H_2_O when applied alone.

### RNA extraction and RNA knockdown

RNA from 2D hiPSC-CMs was extracted with TRIzol® (Thermo Fisher Scientific) according to the manufacturer’s instructions. Concentration, purity and quality of RNA was measured using a NanoDrop® ND-1000 spectrophotometer (Thermo Fisher Scientific).

For knockdown of *PRKG1*, two commercially available siRNAs were used as indicated (Ambion, s11131, s11132). Control reactions in matching concentrations were performed using scramble (*scr*) siRNA (Thermo Fisher Scientific, s7646-48). HiPSC-CMs were plated at a density of 2.4 × 10^5^ cells/well onto 12-well plates coated with Geltrex and cultured as described above. Culture medium was exchanged for 500 µl of OptiMEM (Gibco, 31985070). Simultaneously, siRNAs and Lipofectamine3000 (Thermo Fisher Scientific, L3000008) (4 µL/mL medium) were diluted in OptiMEM and incubated for 5 min. The dilutions were mixed at a 1:1 ratio and incubated for 20 min. Finally, the transfection mixture was added dropwise to the appropriate wells. Four hours after transfection 1 ml of maturation medium was added per well. Medium was changed three days after the transfection and cells were harvested for protein or mRNA analysis after 5 days.

### RT-qPCR analysis

RNA was extracted as described above and complementary DNA was generated from 100-200 ng per sample with iScript cDNA synthesis kit (BioRad, 12192). RT-qPCR analysis for the quantification of *PRKG1A* mRNA was performed with a SYBR Green/Rox qPCR master mix (2X; Thermo Fisher Scientific, K0222) according to the provided manual but with a reduced reaction volume of 10 µl instead of 20 µl and 45 instead of 40 cycles. The primers used for *PRKG1A* (CTCCACAAATGCCAGTCGG; TTATAAGATCCTTGGACCTTTCGG) and the housekeeping gene β-Glucuronidase (*GUSB*; ACGATTGCAGGGTTTCACCA; CACTCTCGTCGGTGACTGTT) were designed with the respective online tool provided by NCBI. Data acquisition was performed in the exponential phase and *PRKG1A* mRNA was quantified relative to *GUSB* with the 2^-Δ^ ^ΔCt^ method.

### RNA sequencing

Total RNA was isolated using the Direct-zol RNA Microprep Kit (Zymo Research) followed by a DNase digestion step. Enrichment of mRNAs with the NEBNext Poly(A) mRNA Magnetic Isolation Module was followed by library preparation using the NEBNext Ultra II RNA Directional Library Prep Kit for Illumina (New England BioLabs). Single read sequencing took place on a NextSeq 2000 System (Illumina), using the corresponding NextSeq 2000 P3 Reagents (50 cycles), with a read length of 72 base pairs. The integrity of the RNA and quality of the library were assessed using a TapeStation 4200 (Agilent). The sequencing data was automatically demultiplexed using the Illumina BCL Convert Software v3.8.2.

FastQ files underwent pre-trimming and post-alignment quality control rounds using FastQC v0.11.7 (https://www.bioinformatics.babraham.ac.uk/projects/fastqc/). Removal of Illumina adapters and low-quality sequences was performed with Trimmomatic v0.38. Reads of length <15 bases, as well as leading and/or trailing bases with quality <3 or no base call, and bases with average quality <15 in a 4-base sliding window were removed. Alignment was performed with HISAT2 v2.1, using the human genome assembly hg38 (Homo sapiens, GRCh38). Mapped reads (primary alignments) were sorted by read name using SAMtools v1.8, and read counts were calculated with HTSeq v0.11.2.

Differential expression was assessed using DESeq2 for all possible comparisons. Raw read counts were filtered to remove genes with less than 10 counts prior to analysis. Here, statistical significance of coefficients was tested using a negative binomial generalized linear model with the Wald test and p-values were adjusted for multiple comparisons following the Benjamini-Hochberg method (*P*_adj_). Genes were considered differentially expressed at *P*_adj_ < 0.05 values.

To provide a biological context, each list of differentially expressed genes (DEGs) was subjected to functional enrichment analysis using the web tool g:GOSt from g:Profiler. All gene ontology (GO) and pathways gene set categories available for Homo sapiens within this tool were retrieved. Gene sets were considered significantly enriched following a hypergeometric test for overrepresentation within annotated genes, corrected for multiple comparisons using the Benjamini-Hochberg method (*P*_adj_<0.05).

Data are expressed as log2 ratio, which represents the difference in log2 mean value between the test and reference groups (=log2[test]−log2[reference]) where the mean of log2(reference) was set to 0.

### Western blot analysis

Cells were harvested in SDS sample buffer (0.03 M Tris pH 8.8, 5 mM EDTA, 30 mM NaF, 3% SDS, 10% Glycerol) supplemented with 2 mM Dithiothreitol (DTT). Before loading the samples onto 4-15 % Mini-Protean TGX precast protein gels (BioRad Laboratories, 4561086), 6X loading buffer (Thermo Fisher Scientific, J61337.AD) was added in the appropriate amount. Afterwards samples were boiled at 95 °C for 5 min and subsequently loaded onto the gel. For gel electrophoresis, voltage was set to 80 V for 10 min and afterwards to 150 V until the desired resolution was achieved. Blotting was performed using a Trans-Blot Turbo Transfer System (BioRad Laboratories) and nitrocellulose transfer packs (BioRad Laboratories, 1704158) with the mixed molecular weight preset. Ponceau S was used to stain for total protein and membranes were blocked using 5% milk powder or 5% bovine serum albumin in 0.1% TBS-T or PUREBlock (Vilber, PU4010500). Primary antibodies (**Table S2**) were incubated over night at 4 °C while secondary antibodies were incubated at room temperature for 90-120 min. Secondary antibody (**Table S2**) signals were captured with ECL solution or appropriate fluorescence excitation and detection on the FUSION FX7 (Vilber). The fluorescently labelled GAPDH680 (Invitrogen, MA5-15738-D680) loading control antibody was incubated for 1 h at room temperature.

### PRKG1 inhibition

For inhibition of PRKG1 the inhibitor RP8 was used at concentrations ranging from 0.0001 µM to 10 μM for 45 min. The concentration of 10 nM was used in the final experiment vs. vehicle (H_2_O).

### cGMP ELISA

HiPSC-CMs were stimulated with 100 nM CNP, 40 µM sac or vehicle diluted in culture medium for 20 min at 37 °C. Afterwards, the medium was removed, and cells were lysed with 500 µl 1 M HCl, that was supplied with the ELISA kit. The ELISA assay was performed according to manufacturer’s instructions (Sigma Aldrich, CG200).

### ANP, BNP and CNP ELISA

Natriuretic peptides were measured in hiPSC-CMs supernatant samples according to the manufacturer’s instructions. We used the ANP Competive ELISA Kit (Cat# EIAANP, Thermo Fisher Scientific), the human NPPB/BNP ELISA Kit (Cat# EHNPPB, Thermo Fisher Scientific) and the human C-type natriuretic peptide/CNP ELISA Kit (Cat# EH136RB, Thermo Fisher Scientific).

### FRET imaging

HiPSC-CMs were seeded on glass cover slips in 6-well plates coated with Geltrex diluted in RPMI as described above. Per well/cover slip 1.95 × 10^5^ cells were seeded. After being cultured as described above, cells were transduced with an adenovirus encoding for the cGi500 FRET sensor at a multiplicity of infection of 30 in culture medium. For imaging purposes, the coverslips were removed from the 6-well plates individually, placed in a specific chamber and washed with FRET buffer (144 mM NaCl, 10 nM HEPES, 1 mM MgCl2, 5 mM KCL, pH 7.2–7.3). Following that, 1 mL of said buffer were put on the coverslip in the chamber and the chamber placed on the microscope. A frame with sufficiently transduced cells was chosen and the recording of CFP/YFP signals was started (one frame per 10 s). Before any compound application, a baseline signal was established. After each compound application the experimenter made sure that the signal was stable. For maximum stimulation of cGMP levels 100 nM CNP/100 µM IBMX were used. The signal recording was performed in ROIs defined by the experimenter prior to starting the experiment alongside with a secondary ROI for the background which was then subtracted from the cell ROI by the applied software.

### Immunofluorescence staining for confocal microscopy and image analysis

For immunofluorescence analysis of two-dimensional cultures, cells were plated, and experiments were carried out on black 96-well plates (Greiner, 655090) which were coated with Geltrex. They were seeded and cultured as described above for about 14 days. After the experiment was conducted, wells were washed with warm PBS once and fixed with Roti-Histofix 4% (Carl Roth, P087.1) for 15 min at 4 °C. Afterwards wells were washed twice with warm PBS and stained with a primary antibody against mouse α-actinin-2 (ACTN2; 1:800; Sigma A7811) over night at 4 °C with gentle shaking. The following day, PBS was used for washing and secondary antibody AF-488 mouse (Thermo Fisher Scientific, A11029) was incubated for 2.5 h at RT. Antibody staining was performed in staining solution (1x PBS, 3% milk powder, 0.1% Triton X-100). Finally, Hoechst 33342 was added for 20 min (1:2500; Invitrogen, H3570) in PBS as a DNA stain. Imaging was carried out using a Zeiss LSM 800 confocal microscope. For cell volume measurements, z-stack imaging was carried out and the volume of each slice was calculated using the area positive for ACTN2 and the distance between consecutive slices. This analysis was only carried out on singular cells and final analysis was performed using Fiji.

### Constructs and enzyme purification for TIRF and immunofluorescence studies

The inserts in pETDuET-1 vector encoding human full-length sfGFP-VASH1-SVBP and its truncated version sfGFP-VASH1(CD+Ct)-SVBP were previously described (8). The pETDuET-1 vector encoding the phosphomimic VASH1 sfGFP-VASH1(7E)-SVBP was obtained by PCR amplification of the above plasmid with overlapping primers encoding substitutions S313E, S323E, S330E, S331E, S337E, S344E. PCR was performed with Phusion DNA polymerase and circularization by In-Fusion HD Cloning kit. Coding regions were verified by DNA sequencing (Genewiz). Protein expression and purification of the three His-tagged VASH1-SVBP complexes were performed as previously described (*22*). Concentration of soluble proteins was measured using Bradford assay with BSA as the standard. All proteins were kept in storage buffer (20 mM Tris-HCl at pH 7.4, 150 mM NaCl, 2 mM DTT) and stored in liquid nitrogen until use.

### Constructs and lentivirus production for expression in cells

For plasmid “EYFP-VASH1-IRES-SVBP-myc-DDK + pLV”, separate PCR amplifications of IRES sequence and cDNAs encoding EYFP, mouse VASH1 (NP_796328), and mouse SVBP (NP_077782) were performed with Phusion DNA polymerase (Thermo Scientific). For EYFP PCR, cDNA encoding a PreScission site was added in the reverse primer. For SVBP PCR, cDNA encoding Myc and 6xHis tags was added in the reverse primer. PCR fragments were introduced in lentiviral pLV vector using In-Fusion HD Cloning kit (Clontech). The resulting construct encodes EYFP-PreScission site-Vash1 and SVBP-myc-6His.

For plasmids “EYFP-V1-7E-IRES-SVBP-myc-DDK + pLV” and “EYFP-V1-7A-IRES-SVBP-myc-DDK + pLV”, the entire “EYFP-V1-IRES-SVBP-myc-DDK + pLV” plasmid was amplified by PCR with overlapping primers. The forward primer encoded the substitutions S323E/A, S333E/A, S334E/A, S340E/A, S341E/A, S347E/A and S354E/A. PCRs were performed with Phusion DNA polymerase and plasmid circularization by In-Fusion HD Cloning kit. Coding regions were verified by DNA sequencing (Genewiz).

For production of lentivirus, stocks of VSV-G pseudotyped viral particles were generated by co-transfection of HEK293T cells (293AAV, Cellbiolabs, Cat. No. AAV-100) using lentiviral packaging plasmids psPAX2 (Addgene plasmid #12260) and pMD2.G (Addgene plasmid #12259) and the particular pLV lentiviral transfer plasmids expressing VASH1 mutants fused to YFP. An IRES site allows expressions of SVBP, myc and FLAG tags. 293AAV cells were cultivated in Dulbecco’s modified Eagle’s medium (DMEM, High Glucose, HEPES, no Phenol Red, FischerScientific, Cat. No. 21063045) supplemented with 10% (v/v) heat-inactivated fetal calf serum, 0.1 mM MEM Non-Essential Amino Acids (NEAA, FischerScientific, Cat. No. 11140050), 100 U/mL penicillin and 100 μg/mL streptomycin (FischerScientific, Cat. No. 15070063). Tissue culture reagents were obtained from Sarstedt. Briefly, 1E+07 293AAV cells were seeded one day before transfection on 15-cm culture dishes and transfected with 3.5 µg packaging plasmid psPAX2, 1.75 µg pMD2.G and 10 µg pLV per plate complexed with Polyethylenimine (PEI MAX Transfection Grade Linear Polyethylenimine Hydrochloride MW 40,000, Polysciences, Cat. No. 24765) at a PEI:DNA ratio (w/w) of 3:1 (*43*). The next day, cell culture medium was exchanged for DMEM/FCS supplemented with 2 mM caffeine. Three days after transfection, the cell culture supernatants were harvested and filtered through 0.45 µm membranes (Sarstedt). After concentration by ultracentrifugation for 2h at 4°C (25,000 rpm, SW32Ti rotor, Beckman Coulter) on a 20% sucrose cushion, the pellets were resuspended in DPBS. The functional titers were determined by transduction of 293AAV cells and quantification by flow cytometry (FACS CantoII, BD Biosciences; FITC Channel) and Lentivirus Titer p24 ELISA Kit Pro (GenScript Cat. No. L00974). The lentivirus particles were further used for transduction of cardiomyocytes. The efficiency of transduction was evaluated by fluorescence microscopy.

### PRKG1A in vitro kinase assays

*In vitro* phosphorylation of VASH1 enzymes was carried out using a method adapted from Banko et al (*23*). sfGFP-VASH1-SVBP (2 µM) proteins were incubated for 1 h at 30 °C in the presence or absence of PRKG1A (10 nM, Sigma Aldrich Merck) in a buffer containing 40 mM Tris-HCl pH 7.5, 100 mM NaCl, 10 mM MgCl_2_, 5% glycerol, 1 mM dithiothreitol and 60 μM ATPγS (Abcam). The kinase reaction was stopped by addition of 20 mM EDTA. Alkylation was then performed by adding 50 mM PNBM (p-nitrobenzyl mesylate, Abcam) for 1 h and stopped with the addition of Laemmli buffer. Phosphorylation of proteins was detected in immunoblot analysis using anti-thiophosphate ester RabMAb (Abcam, ab92570, 1:5000) and secondary anti-rabbit HRP antibody (Jackson ImmunoResearch 711-035-152; 1:10,000).

### Preparation of tubulin and taxol-stabilized microtubules

Brain tubulin purification and its labeling with either biotin or ATTO-565 fluorophore (ATTO-TEC GmbH) were done as previously described (*44*). Tubulin from HeLa cells were produced and purified as previously described (*45*) with some modifications. HeLa S3 cells (CLS Cell Lines Service GmbH) were first cultivated ∼5 days in adherent cultures and then used to inoculate 2 L of suspension culture that were cultured during ∼14 days at 65 rpm, 37 °C, 5% CO_2_ in a New Brunswick S41i incubator (Eppendorf). Cells were then harvested, resuspended in a lysis buffer (BRB8O (80 mM PIPES (piperazine-N, N′-bis (2-ethanesulfonic acid))-KOH at pH 6.8, 1 mM MgCl_2_, 1 mM EGTA) supplemented with 1 mM 2-mercaptoethanol, 1 mM phenylmethylsulphonyl fluoride (PMSF), cOmplete EDTA-free protease inhibitor cocktail tablets (GE Healthcare) and 0.2 % (vol/vol) Triton X-100, and lysed by sonication. Cell lysate was clarified by centrifugation at 150 000g. The supernatant was then subjected to two rounds of polymerization/depolymerization cycles (one in a low molarity buffer containing 0.08 M PIPES, the other one in high-molarity buffer containing 0.5 M PIPES, both with GTP and glycerol), as previously described (*45*). The polymerized microtubules were finally spinned down at 150 000 g for 30 min at 30 °C. The pellet obtained was depolymerized on ice using cold BRB80 (20-30 min), followed by a centrifugation at 200 000 g for 20 min at 4 °C. The supernatant contains exclusively tyrosinated tubulin.

Taxol-stabilized microtubules were prepared as described in (*22*). Microtubules were polymerized using 45 μM tubulin (composed of 65% of HeLa tyrosinated tubulin, 30% biotinylated brain tubulin, and 5% ATTO-565–labeled brain tubulin) in BRB80 supplemented with 1 mM GTP. After microtubule formation, 100 μM Taxol were added and incubation was continued for 30 min. Tyr-microtubules were then centrifuged 10 min at 200,000 x g and resuspended in BRB80 supplemented with 10 μM Taxol.

### TIRF in vitro assay

TIRF imaging assays were performed as previously described (*22*). Taxol-stabilized microtubules enriched in Tyr-tub were perfused into TIRF chambers and incubated at room temperature for 5 min. Unbounded microtubules were removed by three washes with 100 µl of BRB80 supplemented with 1% bovine serum albumin (BSA) and 10 μM taxol. Finally, 30 µl of a solution containing 50 pM of the sfGFPVASH1–SVBP complex in BRB80 (supplemented with 82 µg/mL catalase, 580 µg/mL glucose oxidase, 1 mg/mL glucose, 4 mM DTT, 0.5 mM taxol, 0.01% methylcellulose 1,500 cp) were perfused. The chamber was sealed, and images were recorded within the first 30 min following addition of the assay-mix solution on an inverted microscope (Eclipse Ti, Nikon). The microscope was equipped with a Perfect Focused System, a CFI Apochromat TIRF 100×/1.49 N.A. oil immersion objective (Nikon), a warm stage controller (Linkam Scientific) and a Technicoplast chamber to maintain the temperature, an objective heater (OkoLab), an iLas2 TIRF system (Roper Scientific), and a sCMOS camera (Prime95B, Photometrics) controlled by MetaMorph software (version 7.10.3, Molecular Devices). For dual-view imaging, an OptoSplit II bypass system (Cairn Research) was used as image splitter, and illumination was provided by 488- and 561-nm lasers (150 and 50 mW, respectively). Temperature was maintained at 35 °C for all imaging purposes. Acquisition rate was one frame each 50 ms exposure (in streaming acquisition) during 45 s.

### In vitro detyrosination activity assay by immunofluorescence

VASH1 activity was measured as previously described (*22*). Taxol-stabilized microtubules were perfused in the same perfusion chambers and TIRF experiments conditions as above. After the addition of VASH1-SVBP enzymes (50 pM), chambers were incubated at 37 °C for 30 min followed by three washes with 10 µM taxol and 1% BSA in BRB80 (wash buffer). The incubation with primary custom-made antibodies (rat anti-Tyr-tub (YL1/2, 1:6,000) and rabbit anti-dTyr-tub (1:1,000)) was done for 15 min, followed by three washes with 100 µl of wash buffer. Secondary antibodies (anti-rat coupled to Alexa Fluor 488 (Jackson ImmunoResearch, 712-545-153; 1:500) and anti-rabbit coupled to Cyanine 3 (Jackson ImmunoResearch, 711-165-152; 1:500), both diluted to 1:500) were then added and incubated for 15 min, followed by three washes with 100 µl of wash buffer. Images were obtained immediately after using a LEICA DMI600/ROPER microscope controlled by Metamorph Video software. Immunofluorescence and tracking of single VASH1–SVBP molecules for estimation of activity and binding parameters were analyzed using FIJI software and a homemade plugin, respectively, as in (*22*).

### Statistical analysis

The precise number of samples (N), technical replicates (wells; n), biological replicates (batches of cardiac differentiation; d), and outliers is indicated in the figure legends. Statistical analyses were performed with GraphPad Prism8 or Prism10 (GraphPad Software, Inc, Boston, MA), and data are expressed as mean±SEM. Data were assessed for normality with the Shapiro-Wilk test. For the data that did not pass this test, we applied the Kolmogorov-Smirnov test. Statistical significance for 2-group comparison was performed with the unpaired Student *t-*test. Multiple group comparison was performed with 1-way ANOVA and Dunnett’ multiple comparisons test or with the nonparametric Kruskal-Wallis test and Dunn’s multiple comparisons test. When no equal SDs were assumed, we used the Brown-Forsythe and Welch ANOVA test and Dunnett T3 multiple comparisons test. For experiments with multiple variables (groups and treatments e.g.), 2-way ANOVA was used. The calculated p-values are indicated in the respective figures.

## Acknowledgments

The authors gratefully acknowledge Anika Witten, Anne Dietrich, Nils Koppers and Andreas Huge (Münster University) for RNA preparation and RNA-seq run, Roberta Kurelic (Experimental Cardiology, UKE, Hamburg) for contributing to the FRET assays, Danny Schreier (Clinical Chemistry, UKE, Hamburg) for contributing to ELISA on natriuretic peptides, Kristin Wenzel, Karina Grube and Julia Rüdebusch (Greifswald University) for the experiments in mice, the fluorescence-activated cell sorting core facility (UKE, Hamburg) for cardiomyocyte counting, and Amélie Meyer (Experimental Pharmacology and Toxicology, UKE, Hamburg) for her help in sample preparation. We would also like to thank Friederike Cuello and Thomas Eschenhagen (Experimental Pharmacology and Toxicology, UKE, Hamburg) for fruitful discussion. We warmly thank Christophe Bosc for KinasePhos 2.0 analysis of the VASH1 protein, and him and Béatrice Blot for cloning the VASH1 constructs enabling expression in bacteria and lentiviruses production. We also acknowledge the Photonic Imaging Center of Grenoble Institute Neuroscience (Univ Grenoble Alpes – Inserm U1216), which is part of the ISdV core facility. Some figures are extracted from the PhD thesis of Chadni Sanyal (Grenoble University, July 2024) and Moritz Meyer-Jens (December 2025, Hamburg University).

## Funding

This work was supported fully or in part by the Leducq Foundation (20CVD01) to LC and MJM, the German Centre for Cardiovascular Research (DZHK; Grant B17-014 and main cluster) and the German Ministry of Research Education (BMBF) to LC and SK, the Helmut und Charlotte Kassau Stiftung (Hamburg) to MMJ and LC, and by the Agence National de la Recherche (ANR-20-CE16-0021) to MJM.

## Author contributions

Conceptualization – LC, MJM; Methodology – MMJ, CS, NP, SRR, EK, SaS, IB, MHR; Investigation – MMJ, CS, NP, SRR, EK, SaS, MHR; Visualization – MMJ, CS, NP, SRR, VN, MHR, SaS, MJM, LC; Funding acquisition – MJM, LC; Project administration – LC; Supervision – VN, MF, SK, SaS, MJM, LC; Writing – original draft – MMJ, LC; Writing – review & editing – CS, NP, SRR, MHR, MF, VN, SK, SaS, MJM, LC.

## Competing interests

LC is consultant for Bayer AG. All other authors do not have any conflict of interest.

## Data and materials availability

The RNA-sequencing (RNA-seq) data have been deposited to RNA-seq data sets to National Center for Biotechnology Information (NCBI) Gene Expression Omnibus of the European Nucleotide Archive at European Molecular Biology Laboratory-European Bioinformatics Institute (EMBL-EBI) with the accession number GSE317137.

