## Supplementary Materials for "Neprilysin inhibition reduces microtubule detyrosination in cardiomyocytes through a cGMP-PRKG1-VASH1 axis"

*Meyer-Jens et al.,*

### Supplementary Materials

#### Supplementary Figures

#### Full, uncut Western blot membranes

### Supplementary Figures

Fig. S1: Evaluation of dTyr-tub levels in mice and measurement of concentrations of natriuretic peptides in hiPSC-CM medium.

Fig. S2. GO term pathways enrichment of RNA-sequencing analysis in ET1 vs. vehicle and ET1/sac vs. ET1.

Fig. S3. Evaluation of the cGMP-PRKG1 signaling axis in hiPSC-CMs.

Fig. S4. Generation and characterization of VASH1-KO hiPSC lines.

#### Figure S1

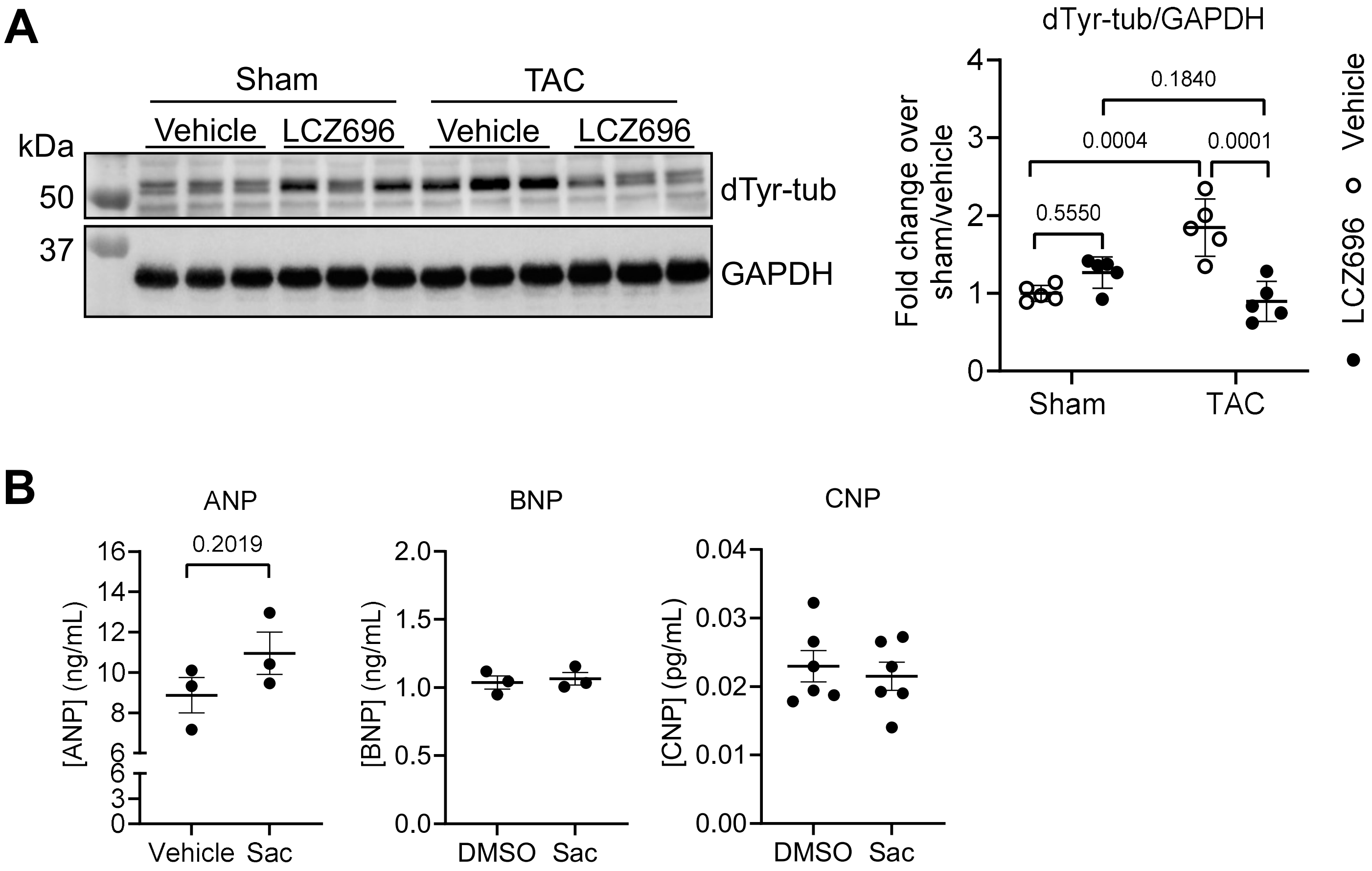


**Fig. S1. Evaluation of dTyr-tub levels in mice and measurement of concentrations of natriuretic peptides in hiPSC-CM medium.** (**A**) Eight-week-old mice were either sham- or TAC-operated and after 2 weeks, they received either a vehicle ($H_20$, 2x 200 µL) or LCZ696 (114 mg/kg body weight/day in a total volume of 2 × 200 µl $H_2O$) per oral gavage for a total duration of six weeks (*16*); representative Western Blot analysis of cytosolic mouse protein extracts stained for dTyr-tub and GAPDH (loading control) and quantification of dTyr-tub/GAPDH (N=5). (**B**) Wild-type hiPSC-CMs were kept in maturation medium for 2 weeks and then treated overnight with vehicle (DMSO, 0.08%) or sac (40 µM in DMSO) for 25 h (ANP and BNP) or overnight (CNP); ELISA analysis of ANP, BNP and CNP concentrations in hiPSC-CM medium (N/d=3/1). Data are expressed as mean±SEM. Statistical significance was assessed with a 2-way ANOVA and Tukey's multiple comparisons test (panel A) or unpaired Student's t-test (panel B). Abbreviations: ANP, atrial natriuretic peptide; BNP, brain natriuretic peptide; CNP, C-type natriuretic peptide; dTyr-tub, detyrosinated tubulin; GAPDH, glyceraldehyde-3-phosphate dehydrogenase; LCZ696, sacubitril/valsartan; MW, molecular weight marker; N/d, number of samples/differentiation; sac, sacubitrilat; TAC, transverse aortic constriction.

### Figure S2

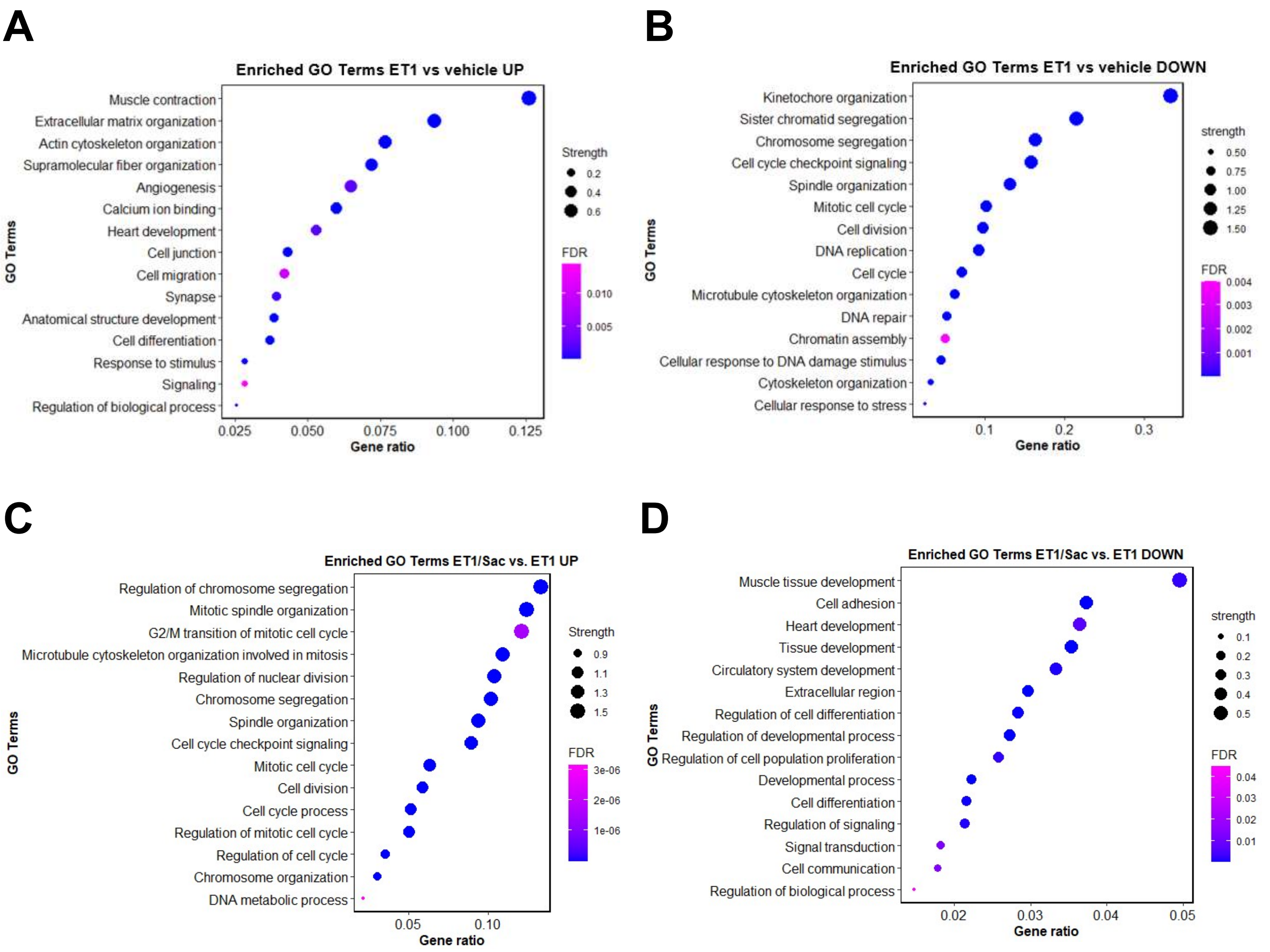


**Fig. S2. GO term pathways enrichment of RNA-sequencing analysis in ET1 vs. vehicle and ET1/sac vs. ET1.** RNA-sequencing was performed in wild-type hiPSC-CMs treated with vehicle (DMSO; 0.08%; N=2), ET1 (100 nM in 0.08% DMSO; N=3) or ET1/sac (100 nM/40 μM in 0.08% DMSO; N=3) for 72 h. (**A**) GO term pathway enrichment of mRNAs with a Log2 ratio > 1 and $P_{adj}$<0.05 in ET1 vs. vehicle. (**B**) GO term pathway enrichment of mRNAs with a Log2 ratio < 1 and $P_{adj}$<0.05 in ET1 vs. vehicle. (**C**) GO term pathway enrichment of mRNAs with a Log2 ratio > 1 and $P_{adj}$<0.05 in ET1/sac vs. ET1. (**D**) GO term pathway enrichment of mRNAs with a Log2 ratio < 1 and $P_{adj}$<0.05 in ET1/sac vs. ET1. The *x* axis shows the gene ratio, the dot size is proportional to the number of gene counts (strength), and the heatmap color shows the extent of $P_{adj}$ values (FDR) from the lowest (purple) to the highest (blue).

**Figure S3**

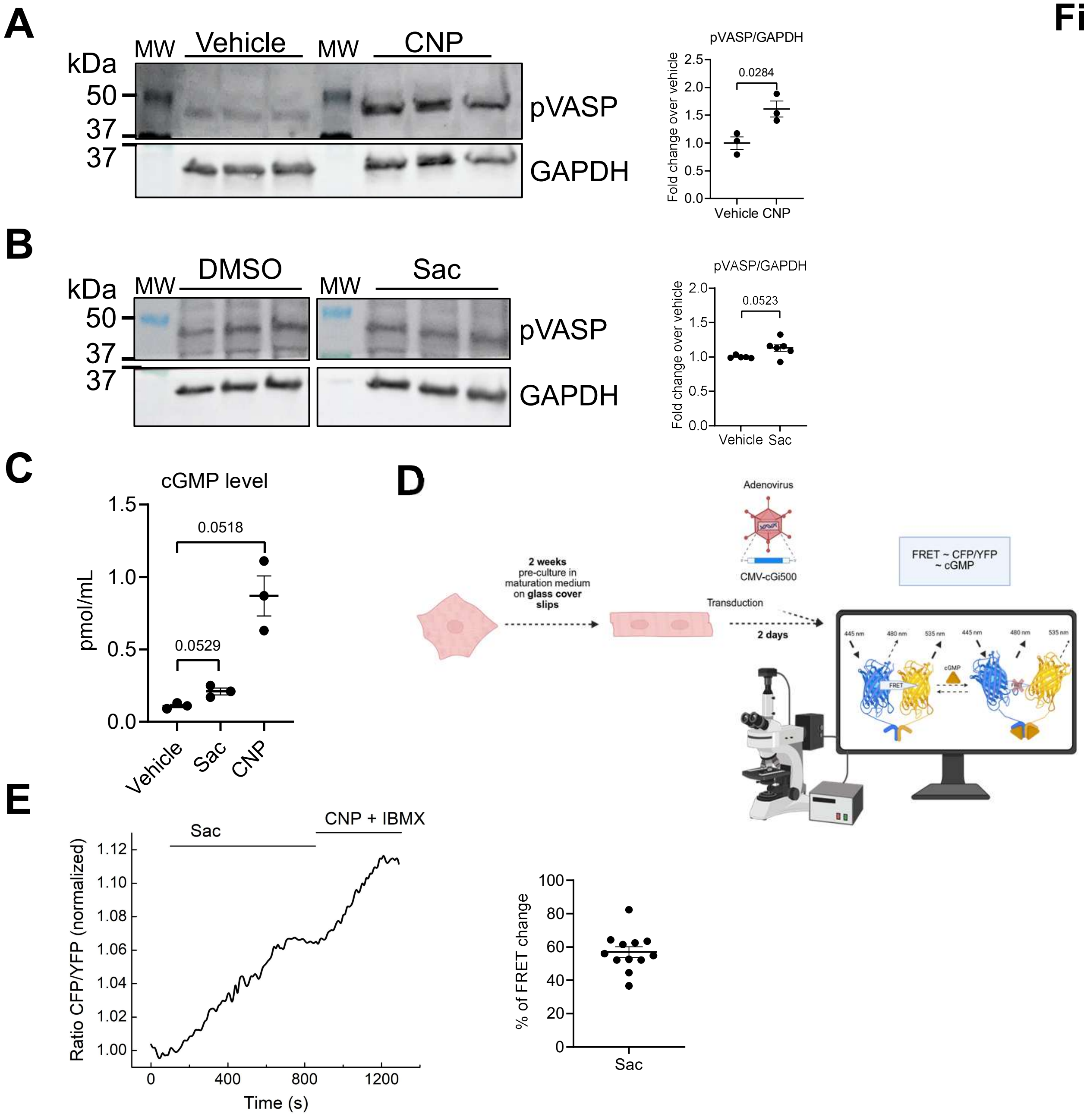


**Fig. S3. Evaluation of the cGMP-PRKG1 signaling axis in hiPSC-CMs. (A)** Wild-type hiPSC-CMs were maintained for 2 weeks in maturation medium and treated with vehicle ($H_2O$) or CNP (100 nM) for 20 min; Western blot of crude hiPSC-CM protein extracts stained for pVASP and GAPDH (loading control) and quantification of pVASP/GAPDH (N/d=3/1). **(B)** Wild-type hiPSC-CMs were treated with vehicle (DMSO, 0.08%) or sac (40 µM in 0.08% DMS0) overnight; Western blot of crude hiPSC-CM protein extracts stained for pVASP and GAPDH (loading control) and quantification of pVASP/GAPDH (N/d=6/1); 1 outlier (robust regression and outlier removal [ROUT] 1%) was removed from the vehicle group. **(C)** ELISA assay for cGMP of wild-type hiPSC-CMs treated for 20 min with vehicle (DMSO 0.08%), sac (40 µM in 0.08% DMSO) or CNP (100 nM in 0.08% DMSO); each dot (N=3) consists of two pooled wells from a 6-well plate. **(D)** Protocol for FRET measurements: wild-type hiPSC-CMs were maintained for 2 weeks in maturation medium and were then transduced for 2 days with the cGi500 FRET adenoviral sensor, consisting of the tandem cGMP-binding domains of PRKG1 sandwiched between donor (CFP) and acceptor (YFP) fluorophores. When CFP is excited and CFP and YFP emission are recorded simultaneously, the emission ratio CFP/YFP of cGi500 can be used to determine the concentration of cGMP; image created with BioRender.com. **(E)** Representative cGMP-FRET measurement in a single cGi500-transduced hiPSC-CM and quantification of percentage of FRET change in 12 hiPSC-CMs. CFP/YFP ratio was measured in baseline condition (DMSO 0.08%), after the application of 40 µM sac (in 0.08% DMSO), and after the application of 100 nM CNP and 100 µM IBMX for a maximal response. Data are expressed as mean±SEM. Statistical significance was assessed with the unpaired Student's *t*-test (panels A and B), or Brown-Forsythe and Welch Anova, followed by Dunnett's T3 multiple comparisons test (panel C). Abbreviations: cGMP, cyclic guanosine 3',5' monophosphate; CNP, C-type natriuretic peptide; ELISA, enzyme-linked immunosorbent assay; FRET, Förster resonance electron transfer; GAPDH, glyceraldehyde-3-phosphate dehydrogenase; hiPSC-CMs, human induced pluripotent stem cell-derived cardiomyocytes; IBMX, 3-isobutyl-1-methylxanthin; MW, molecular weight marker; N/d, number of wells/differentiations; PRKG1, cGMP-dependent protein kinase 1; pVASP, phosphorylated vasodilator-stimulated phosphoprotein at serine 239; sac, sacubitrilat.

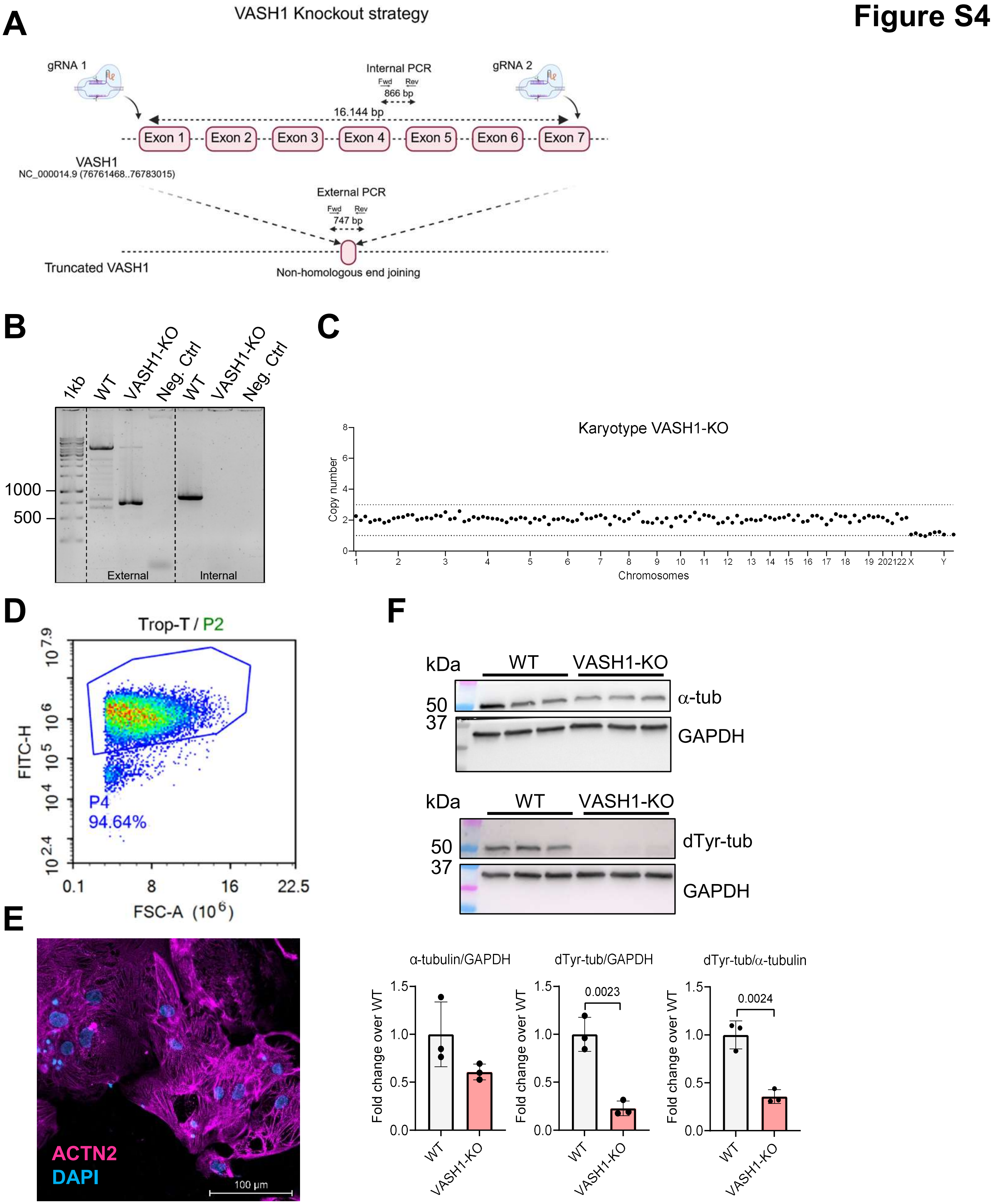


**Fig. S4. Generation and characterization of VASH1-KO hiPSC lines.** (**A**) KO strategy for VASH1. Two gRNAs were used to excise a targeted fragment of 16,144 bp, removing almost the complete coding sequence of VASH1 after non-homologous end-joining. (**B**) KO validation with primers flanking the deletion site (External) and primers inside the targeted locus (Internal). (**C**) NanoString karyotyping of the VASH1-KO hiPSCs. (**D**) FACS analysis of VASH1-KO hiPSC-cardiomyocytes after differentiation indicating successful differentiation with 94.64 % of cells expressing cardiac troponin T. (**E**) Confocal imaging of VASH1-KO hiPSC-cardiomyocytes stained for ACTN2 (magenta) and DAPI (nuclei; blue) shows hiPSC-cardiomyocytes with sarcomeric striations. (**F**) Western blot analysis of hiPSC-cardiomyocytes (7 days in culture). VASH1-KO hiPSC-cardiomyocytes show significantly reduced levels of dTyr-tub normalized to GAPDH or α-tubulin. Statistical significance was assessed with the unpaired Student's *t*-test (panel F). Abbreviations: ACTN2, α-actinin 2; α-tub, α-tubulin; bp, base pairs; dTyr, detyrosinated tubulin; gRNA, single guide RNA; KO, knock-out; TropT, cardiac troponin T; VASH1, vasohibin 1; WT, wild-type.

### Full, uncut Western blot membranes

Uncut membranes Fig. 1

1C)

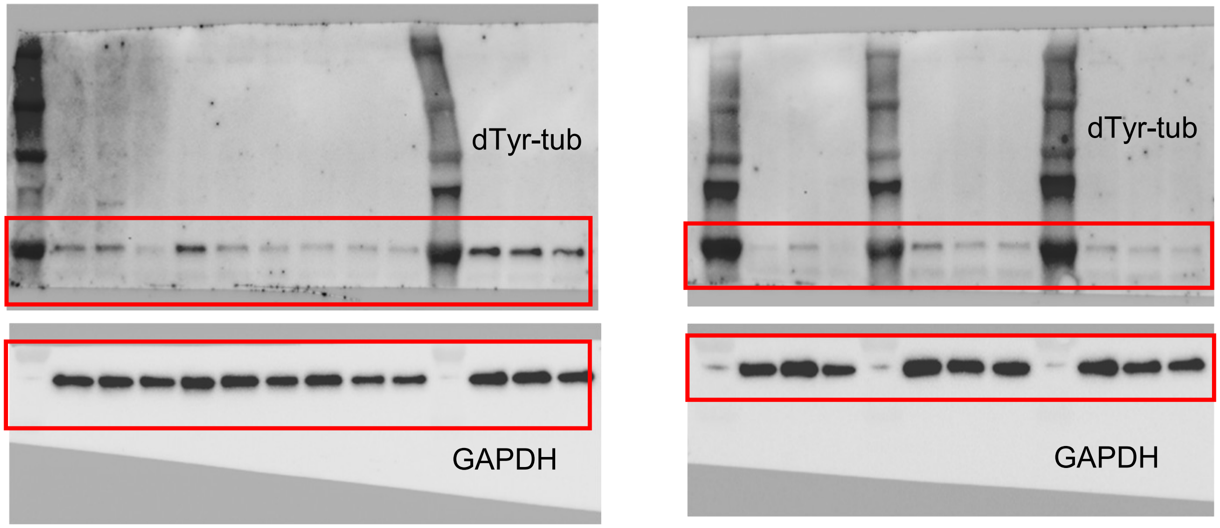


1D)

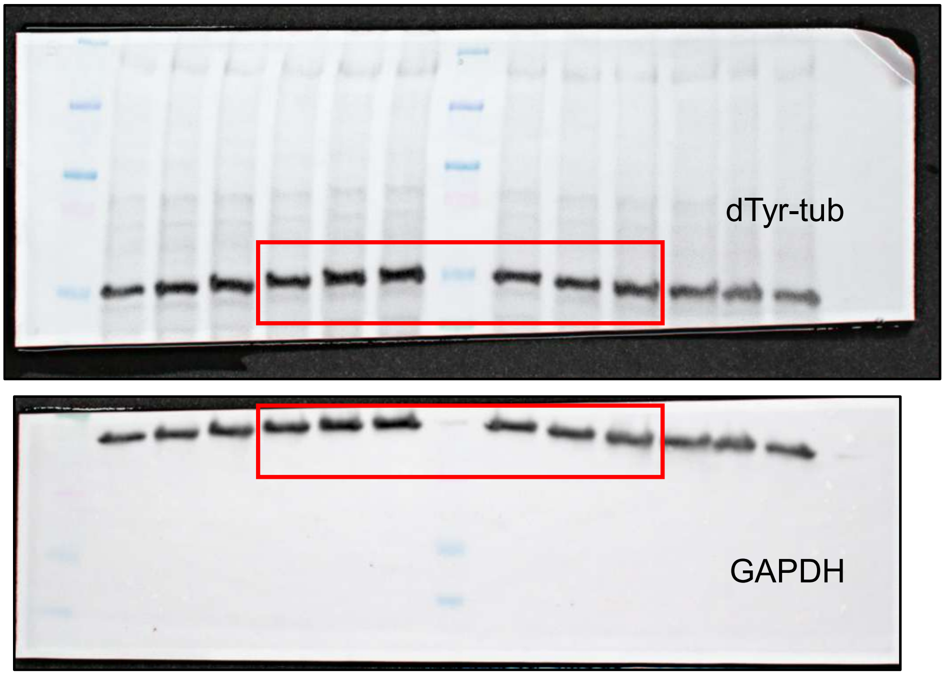

### Uncut membranes Fig. 3

3B)

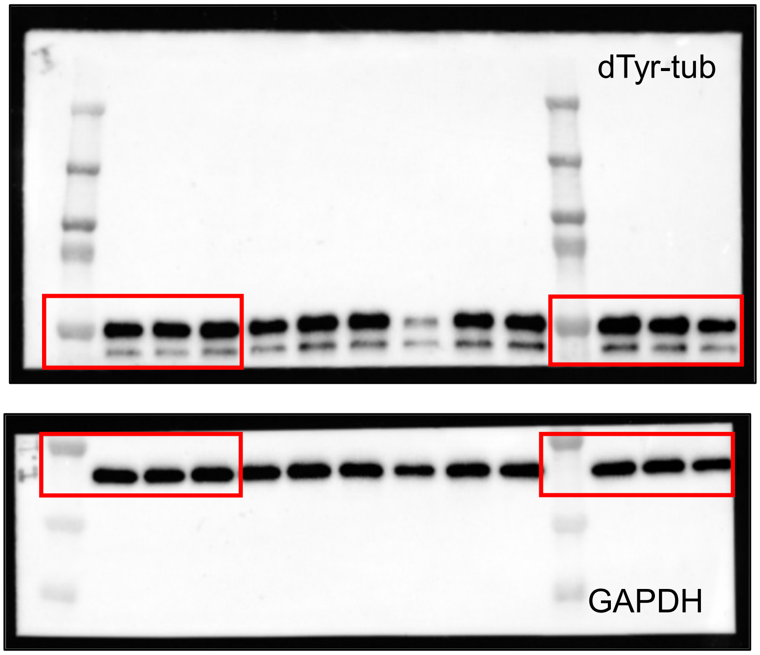


3D)

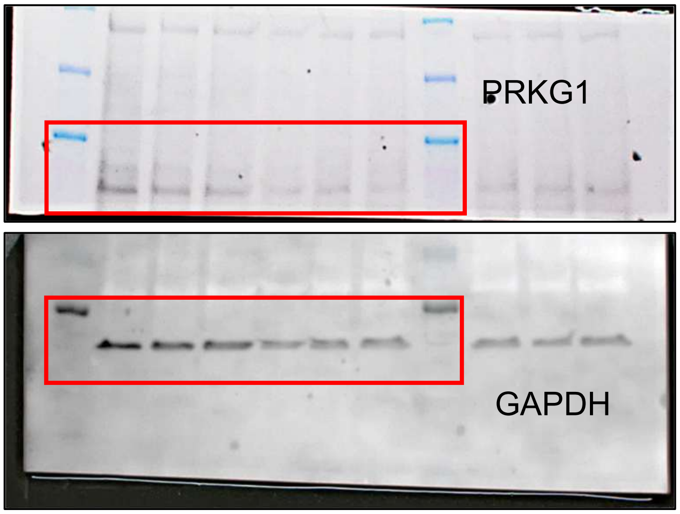


3E)

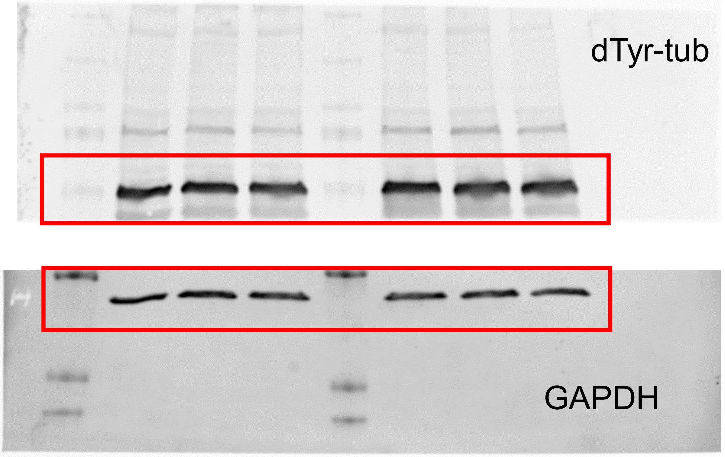

### Uncut membranes Fig. 4

**4B)**

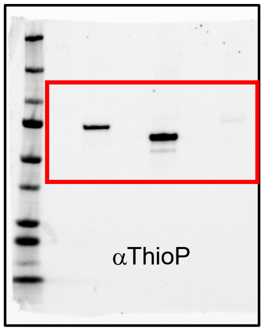


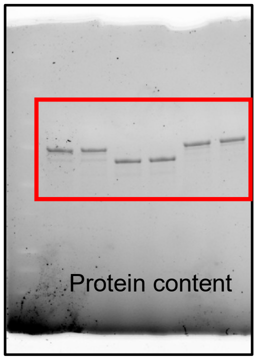

### Uncut membranes Fig. 5

**5C)**

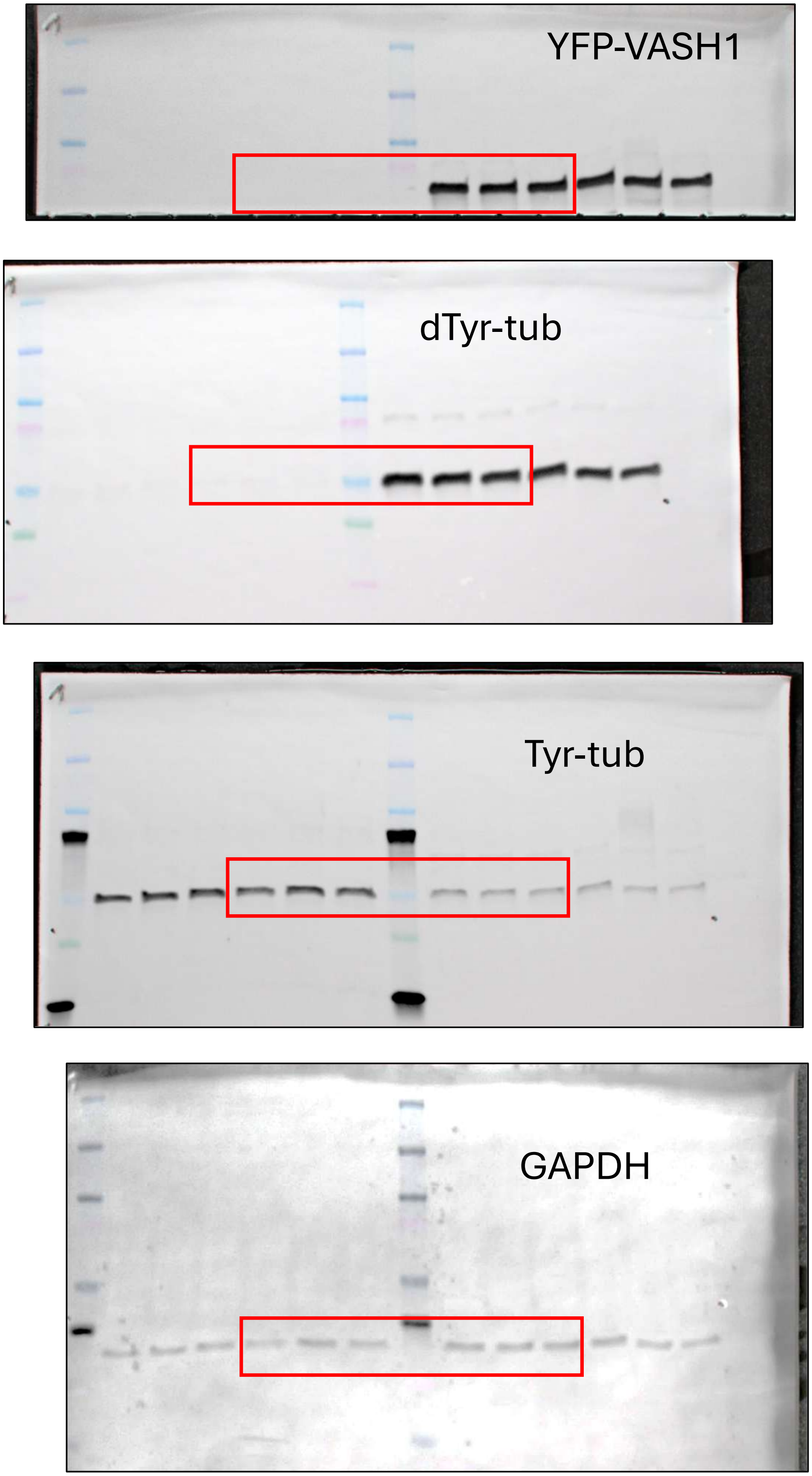

### Uncut membranes Fig. 5

**5D)**

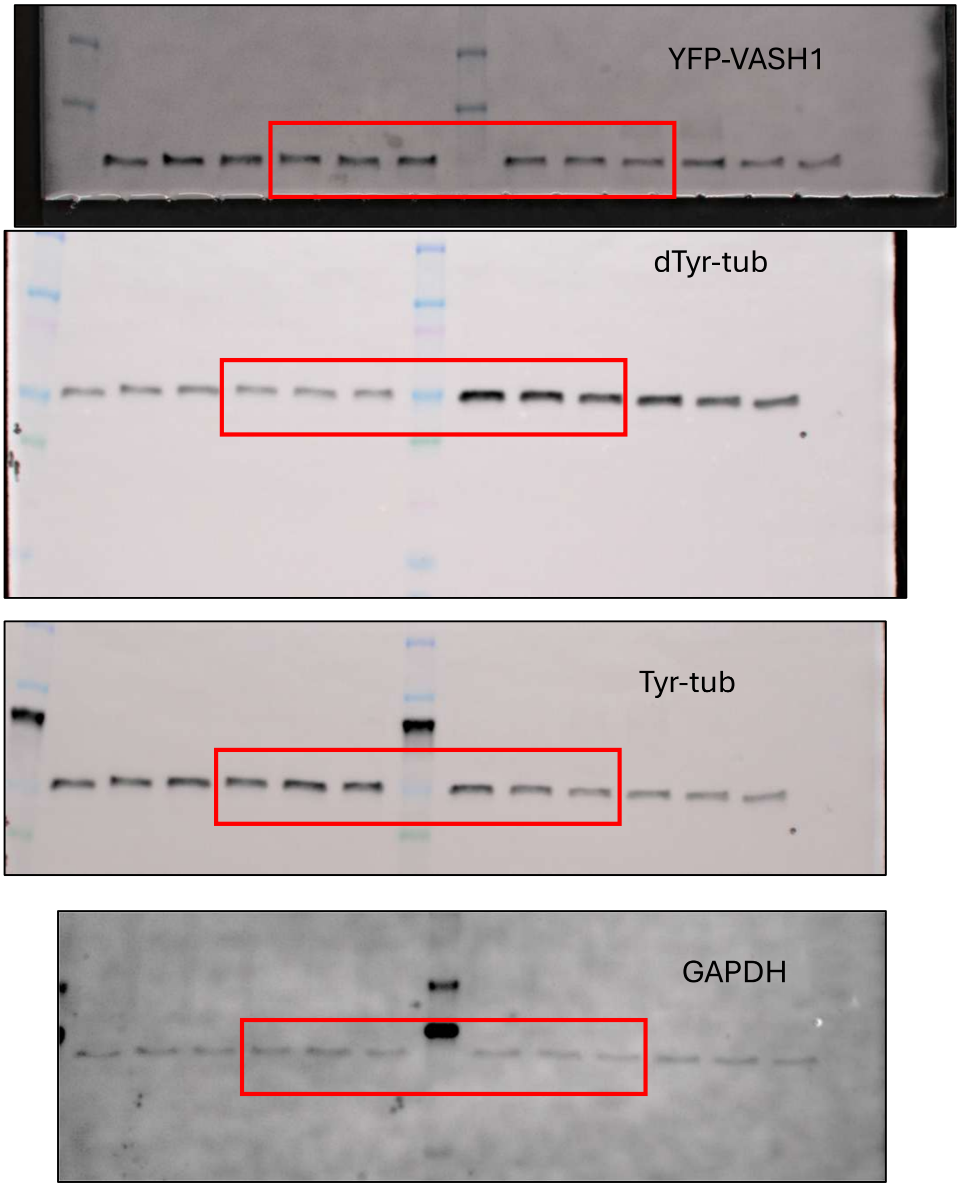
